# Loss of PI5P4Kα slows the progression of a *Pten* mutant basal cell model of prostate cancer

**DOI:** 10.1101/2024.08.12.607541

**Authors:** Joanna Triscott, Marika Lehner, Andrej Benjak, Matthias Reist, Brooke M. Emerling, Charlotte K.Y. Ng, Simone de Brot, Mark A. Rubin

## Abstract

While early prostate cancer (PCa) depends on the androgen receptor (AR) signaling pathway, which is predominant in luminal cells, there is much to be understood about the contribution of epithelial basal cells in cancer progression. Herein, we observe cell-type specific differences in the importance of the metabolic enzyme phosphatidylinositol 5-phosphate 4-kinase alpha (PI5P4Kα; gene name *PIP4K2A*) in the prostate epithelium. We report the development of a basal-cell-specific genetically engineered mouse model (GEMM) targeting *Pip4k2a* alone or in combination with the tumor suppressor phosphatase and tensin homolog (*Pten*). PI5P4Kα is enriched in basal cells, and no major histopathological changes were detectable following gene deletion. Notably, the combined loss of *Pip4k2a* slowed the development of *Pten* mutant mouse prostatic intraepithelial neoplasia (mPIN). Through the inclusion of a lineage tracing reporter, we utilize single-cell RNA sequencing to evaluate changes resulting from *in vivo* downregulation of *Pip4k2a* and characterize cell populations influenced in the established Probasin-Cre and Cytokeratin 5 (CK5)- Cre driven GEMMs. Transcriptomic pathway analysis points towards the disruption of lipid metabolism as a mechanism for reduced tumor progression. This was functionally supported by shifts of carnitine lipids in LNCaP PCa cells treated with *siPIP4K2A*. Overall, these data nominate PI5P4Kα as a target for PTEN mutant PCa.

**One Sentence Summary:** Loss of PI5P4Kα slows cancer progression in prostate basal cells.

## INTRODUCTION

Epithelial lineage diversity enables prostate cancer (PCa) to escape androgen receptor (AR) targeted therapies. The adult prostate epithelium consists of luminal, basal, neuroendocrine, and basal-intermediate cells (*1*, *2*). Initially, PCa depends on AR signaling, which is predominant in luminal cells. The development of castrate-resistant PCa (CRPC) and AR-negative CRPC leads to dysregulated cell lineage plasticity to AR indifferent cells. Clues to how PCa adapts to AR-targeted therapies may be revealed by investigating the disease progression of non-luminal cell populations.

Phosphatase and tensin homolog (*PTEN*) is the most frequently mutated tumor suppressor in human PCa, and its genomic alterations generally occur early in the timeline of progression (*3*). In murine genetic models, PTEN can be deleted in luminal or basal cells to generate PCa (*4*, *5*). Although driven by the same mutation, tumor characteristics of different cells-of-origin show unique characteristics. Specifically, lineage-tracing studies suggest cytokeratin 5 (CK5) positive basal cells contain populations of multipotent stem cells that -under normal conditions-can rapidly regenerate the luminal cell compartment. When transformed, the highly plastic basal cell compartment is believed to drive basal to luminal differentiation that feeds cancer pathology (*1*, *4*, *6*, *7*). In human transcriptomic analysis, basal-cell gene expression profiles are enriched in advanced, metastatic CRPC (*8*, *9*). However, questions remain about how *PTEN* deletion in basal cells develops into PCa and which pathways are critical for its molecular orchestration.

Previous work from our group discovered a relationship between low AR signaling and the alpha isoform (gene name *PIP4K2A*) of the phosphatidylinositol 5-phosphate 4-kinases (PI5P4Ks) and nominated it to have a role in supporting CRPC development (*10*). The enzymatic function of PI5P4K is to phosphorylate the fourth position of the phosphoinositide PI-5-P to produce PI-4,5-P_2_ on membranes of intracellular organelles (*11*, *12*). Known locations such as the lysosome, peroxisomes, and endosomes highlight the function of PI5P4K in regulating stress metabolism and the mammalian target of the rapamycin complex 1 (mTORC1) signaling pathway. In addition, PI5P4K is essential for lipid trafficking and beta-oxidation by facilitating inter-organelle signaling (*10*, *13–15*). In PCa, *PIP4K2A* targeted hairpin systems are efficacious in reducing *in vitro* cell line growth, and luminal-cell targeted deletion has no evidence of morphological abnormalities in the non-castrate setting. Importantly, PI5P4K isoforms are druggable targets receiving attention for oncology drug development (*16–18*). While showing early promise in some cancer types, much is to be understood about the phosphatidylinositol (PI) networks, specifically in PCa biology. For example, while inhibition of PI5P4K may be efficacious for *TP53* mutant breast cancer (*19*), PI5P4Kα is suggested to function as a tumor suppressor in glioblastoma multiforme (GBM) (*20*). Thereby, we experimentally delve deeper into the role of PI5P4Kα in non-luminal biology and combination with *PTEN* deletion.

One of the challenges in studying PI5P4K is how its variable function and expression can change in the context of different cell types. Although shown to regulate critical lysosome signaling events during androgen depletion stress, *PIP4K2A* expression is relatively low in bulk expression analysis of prostate when compared to other tissue types (*10*). We utilize single-cell RNA-sequencing (scRNA-Seq) of enriched basal cell populations to measure how PI5P4K isoform expression varies in the spectrum of transformed PCa cell populations *in vivo*. Also, we demonstrate how deletion of *PIP4K2A* influences the occurrence of mouse prostatic intraepithelial neoplasia (mPIN) at the single-cell resolution. The use of scRNA-Seq methodology has established an early foothold in PCa research in recent years (*2*, *21*, *22*). Expanding on prostate cell atlas datasets, we focus on enriched cell populations using fluorescent marker selection of Cre recombinase activity (*23*, *24*). This approach sheds light on how mutated cells transform into heterogeneous tumor populations.

We aim to understand how the specialized biology of the prostate epithelial basal cell contributes to poor disease outcome. Herein, we use *in vivo* mouse models to characterize the outcome of deleting PI5P4Kα in basal cells with and without co-deletion of the tumor suppressor, *Pten.* In addition, scRNA-Seq analysis reveals the molecular consequence of altering PI5P4Kα and links impaired lipid metabolism with delayed mPIN development. This study uncovers nuances between epithelial cell types and overlays these findings in the context of cancer development. Basal-like subtypes of cancer are often associated with worse outcomes. Establishing a clear understanding of how basal cells specifically contribute to cancer phenotypes will aid in answering questions of PCa heterogeneity and progression.

## RESULTS

### 1. PI5P4Kα has diverse expression in prostate epithelium

The roles of PI5P4K are known to vary between tissue types, and recently, methods have been optimized to consider cell-type-specific differences within native tissues. We initially characterized how PI5P4Kα transcript (*PIP4K2A*) expression compares between basal (BE) and luminal (LE) epithelial prostate cells. A scRNA-Seq dataset of 28,702 human prostate cells was queried for multiple annotated cell types, including BE, LE, club, and hillock (annotations previously described (*25*) (**Fig. 1A**). *PIP4K2A* is a relatively low abundance transcript that is frequently missed in low-coverage sequencing reads, but positive cells were detected in BE (259/424) and LE (61/424) populations (**Fig. 1B and S1, A and B**). In correlative data in bulk RNASeq data confirms an enrichment in BE signatures genes compared to LE genes in human PCa (**Fig. S1C**) (*26*). *PIP4K2A* may have important functions in multiple prostate epithelial cell types.

**Fig. 1.**
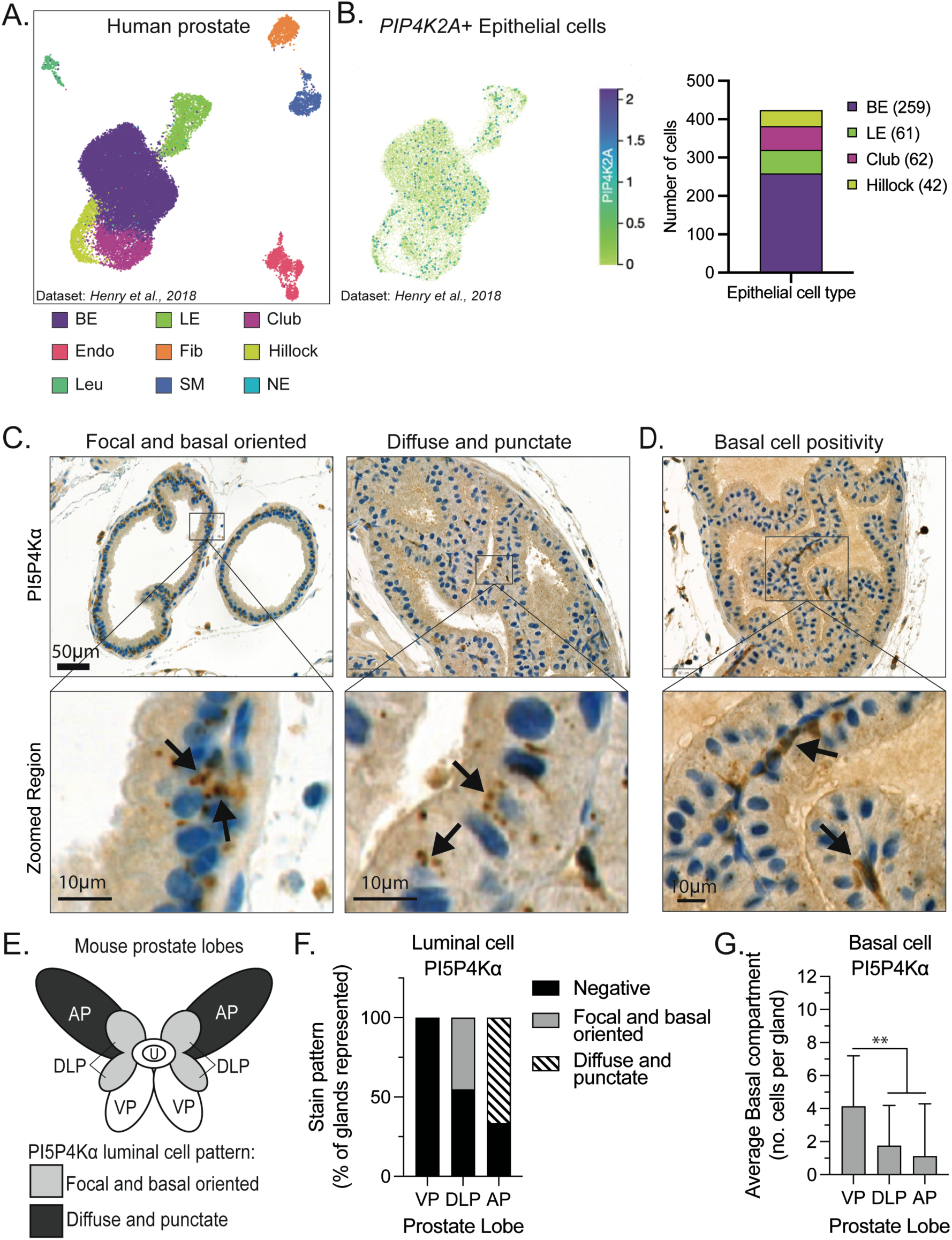
Cell type expression patterns of PI5P4Kα. (**A**) A human adult prostate single cell RNA-Seq (scRNA-Seq) dataset is annotated for basal epithelium (BE), luminal epithelium (LE), club, endothelia (Endo), fibroblasts (Fib), hillock, leukocytes (Leu), smooth muscle (SM), and neuroendocrine (NE) populations (*25*, *42*) (**B**) Populations are subset for epithelial clusters (BE, LE, club, hillock) and 424 epithelial cells are positive detection of *PIP4K2A* transcript. Adult C57BL/6 mouse prostates stained for PI5P4Kα with immunohistochemistry (IHC) and found to express in three patterns: (**C**) focal and basal oriented (FoB), diffuse and punctate (DiP), and (**D**) basal epithelial layer (zoomed scale bars = 10μm). (**E**) Graphical summary of PI5P4Kα luminal cell patterns based on mouse prostate lobe localization. Anterior prostate (AP), dorsal-lateral prostate (DLP), and ventral prostate (VP) are included relative to the urethra (U). PI5P4Kα stain patterns are numerically summarized based on the prostate lobe for (**F**) luminal patterns and as (**G**) the number of positive basal cells per gland. *t* test values: n.s., not significant (*p*> 0.05), * *p* <0.05, ** *p* <0.01, *** *p* < 0.001.

Next, we sought to characterize the protein expression patterns of PI5P4Kα in the adult mouse prostate. Previous work has colocalized PI5P4Kα with organelle markers of the peroxisome and lysosome, however, PI5P4Kα might have additional subcellular localization and functions (*10*, *15*). We classified three distinct patterns using immunohistochemistry (IHC) (**Fig. 1, C and D**). Two patterns occur in the luminal compartment: focal and basal oriented (FoB) or diffuse and punctate (DiP). The third pattern is intermittent basal epithelium staining that shows full cytoplasmic positivity. The occurrence of the luminal patterns are dictated by specific prostate lobes. Whereas the DiP pattern is observed in the anterior prostate lobe, the FoB is predominately in the dorsal-lateral lobes (**Fig. 1E**). In contrast, basal cell PI5P4Kα positivity is present in all mouse prostate lobes, equal between proximal and distal lobe localization, and averaging between 2-4 strong positive cell per gland/ section (**Fig. 1, D and G**). This diversity of protein localization suggests PI5P4Kα may have variable functions in different cell types of epithelial tissues.

### 2. PI5P4Kα is enriched in prostate basal cells

Evidence in **Fig. 1D** suggests that PI5P4Kα is enriched in basal cells; therefore, we measured the protein content of this cell population. Basal cells were labeled *in vivo* using a tamoxifen-inducible cytokeratin 5 (Krt5)-specific Cre recombinase system paired with a *Rosa26eYFP* reporter gene (**Fig. 2A**). This genetically engineered mouse model (GEMM) is referred to as CK5-eYFP. Specific activation was confirmed with recombination rates between 50-60% estimated with eYFP and CK5 immunofluorescent (IF) staining colocalization (**Fig. 2B**). Mouse prostate organoids were generated from animals two months following tamoxifen induction. Organoids included bulk epithelial cell populations with eYFP+ CK5 expressing basal cells (**Fig. 2C**). Subsequent flow cytometry of *in vitro* cultures purified CK5-eYFP lines representative of epithelial basal cells. The CK5-eYFP cells had elevated CK5 and PI5P4Kα protein compared to bulk prostate cultures (**Fig. 2D**).

**Fig. 2.**
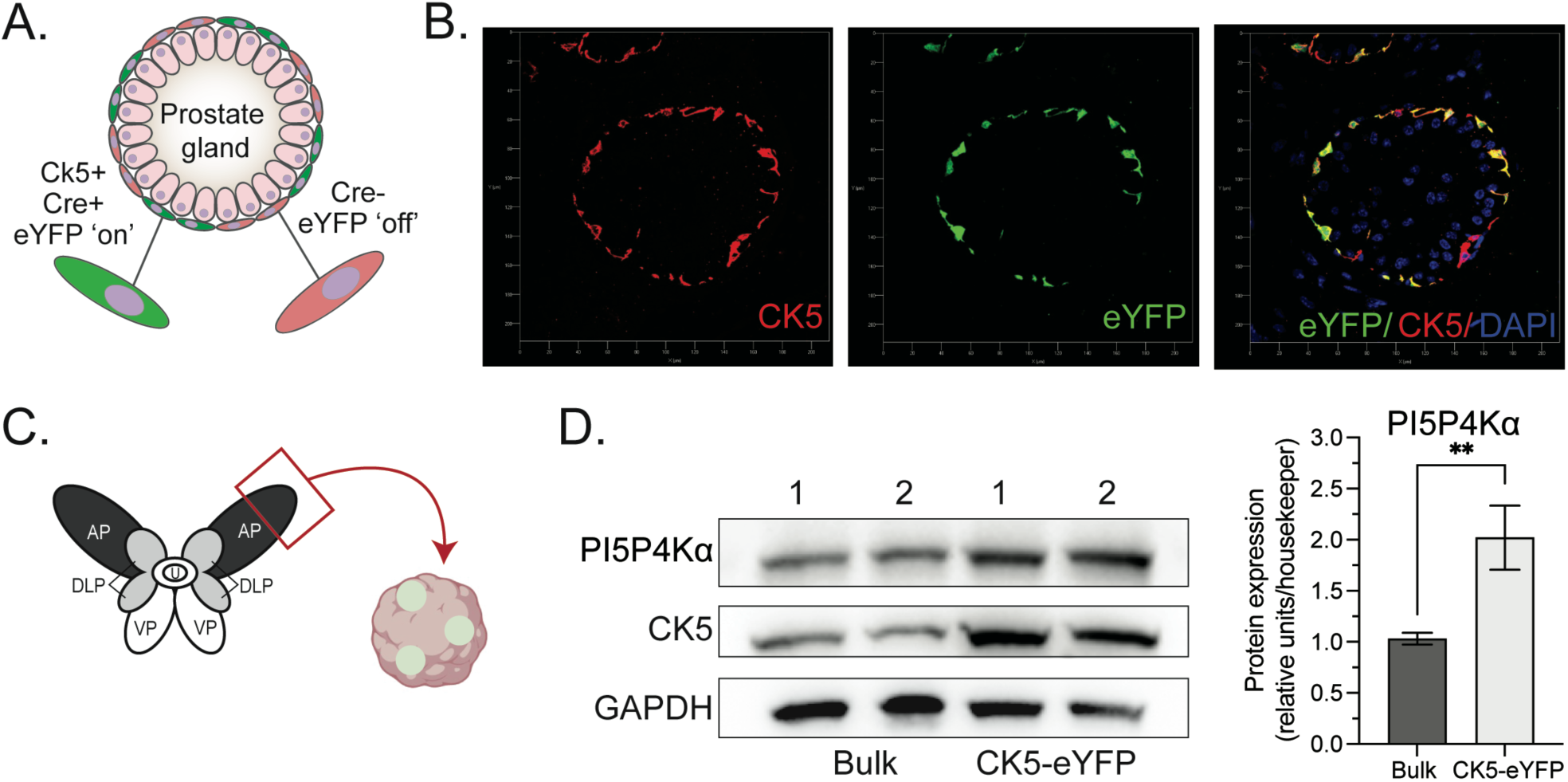
PI5P4Kα is enriched in prostate basal cells. (**A**) Graphical summary of basal cell lineage-marked phenotype of tamoxifen-inducible *Krt5-Cre^ERT2^; R26eYFP* (CK5-eYFP) genetically modified mouse model (GEMM). Fluorescent eYFP protein is expressed in cells where Cre is activated. (**B**) Z-stacked immunofluorescent confirmation of eYFP activation in CK5+ basal prostate cells following. (**C**) 3D organoid cultures generated from the prostate anterior prostate (AP) lobe two months after eYFP activation in the CK5-eYFP model. (**D**) PI5P4Kα and CK5 protein expression in mouse organoid cultures from two biological replicates enhanced following fluorescently activated cell sorting (FACs) of eYFP labeled cells. Protein is quantified from 3 independent experiments relative to housekeeping controls (2.02 fold increase, t.test, pval = 2.9E-03). *t* test values: n.s., not significant (*p*> 0.05), * *p* <0.05, ** *p* <0.01, *** *p* < 0.001.

### 3. Single-cell analysis identifies cell populations influenced by basal and luminal GEMMs

Expression patterns of PI5P4Kα appear to vary based on subpopulations of epithelial cells. In a previous study, we targeted *Pip4k2a* in a luminal cell-specific GEMM (*10*). Before considering the development of new *Pip4k2a-*targeted GEMMs, we wanted to characterize the prostate cell populations impacted by the activation of popular Cre recombinase systems. We implemented a version of scRNA-Seq that uses flow cytometry-based selection of eYFP+ cells from our luminal and basal *in vivo* systems (*24*). The luminal model eYFP+ selected cells were collected from 10-week-old male prostates with a Probasin (PB) -Cre (PB-Cre) recombinase system. We compared this to our CK5-eYFP basal-cell GEMM, the inducible CK5-Cre system, collected two weeks after tamoxifen induction (**Fig. 3A**). Two animals per group and 1500 individual cells were sequenced following FACs selection. Of these, 1042 luminal and 1292 basal model cells passed QC filtering for analysis (**fig. S3A**). Intrinsic cell differences were noted in QC evaluation as luminal model samples had a relatively higher percentage of mitochondrial genes, and basal model samples had more cycling cells (**fig. S5, B and C**).

**Fig. 3.**
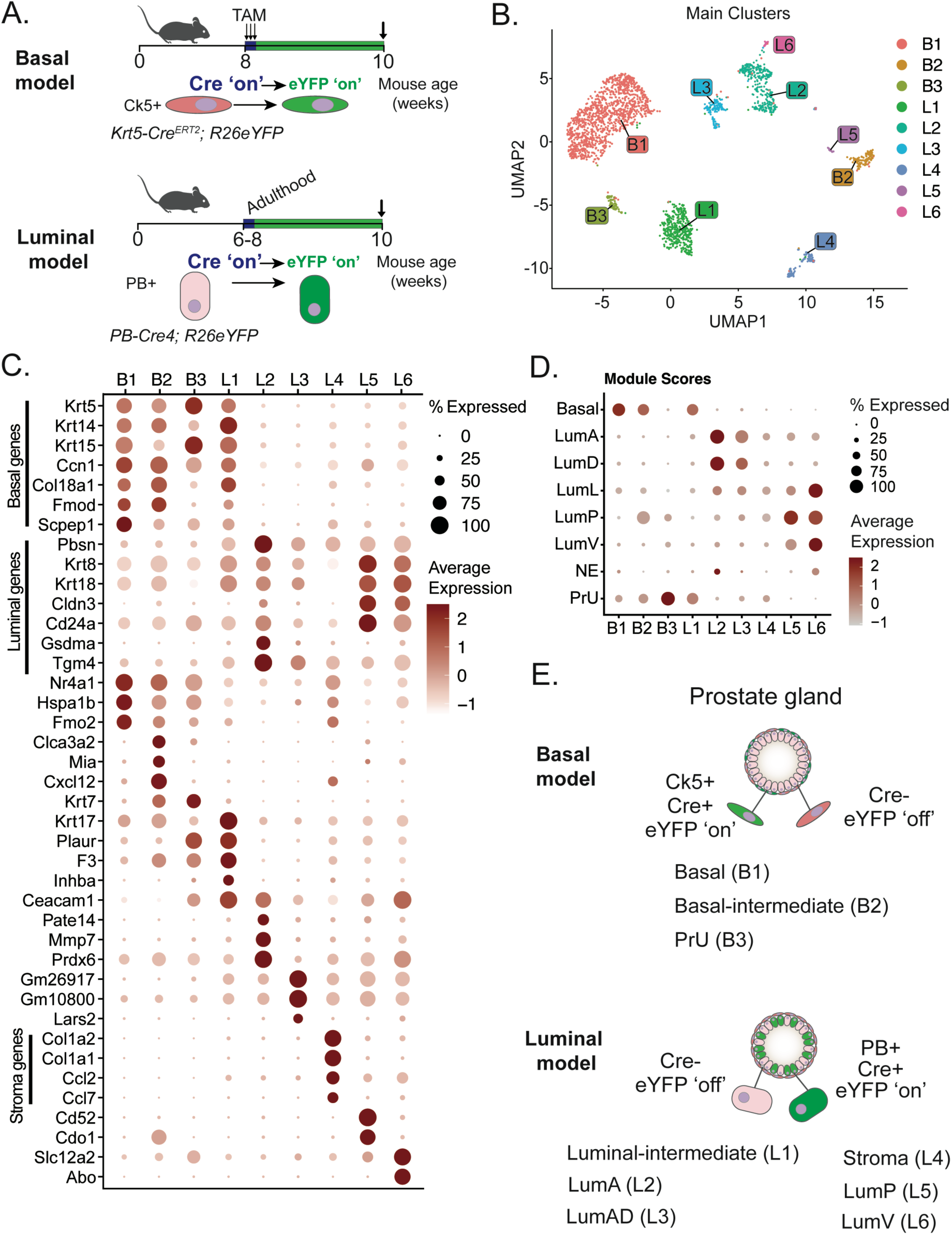
Single-cell RNA sequencing of lineage marked GEMMs. Mouse cells selected from digested *in vivo* prostate tissue were analyzed using the SORT-seq RNA-Seq platform. (**A**) Graphical summary of a probasin (PB) based luminal model and cytokeratin 5 (CK5) based basal model (CK5-eYFP; also see **Fig. 2)**. Cre activation occurs by 8 weeks of age, and eYFP+ cells were collected with FACs two weeks later for both models. (**B**) Uniform Manifold Approximation and Projection (UMAP) clustering of major cell types identified from GEMMs. B1, B2, and B3 are clusters from the CK5-eYFP basal model. L1 to L6 are clusters with origin from the luminal model. (**C**) Clusters have varying levels of canonical basal markers (*Krt5* and *Trp63*) and luminal markers (*Pbsn* and *Krt8*) that have expression associated with expected lineage groups. (**D**) Module scores from marker genes from *Crowley et al.* (see **table S3** gene list) mouse single cell atlas in comparison with targeted scRNA-Seq clusters from this study. (**E**) Summary of predicted cell types that are identified from lineage tracked basal and luminal GEMMs.

Using the graph-based cell clustering method from the Seurat package (*27*), three main clusters were identified from the basal model cells: B1, B2, and B3 (**Fig. 3B**), with most cells falling into a uniform B1 cluster. More diversity of clustering occurred in the PB-Cre model cells, which were divided into four main clusters (L1, L2, L3, and L4) and two additional small clusters (L5 and L6) (**Fig. 3B**). As expected, most cells from the basal and luminal models clustered separately (**fig. S3C**). To confirm the lineage identity of the clusters, known marker genes for basal and luminal cell lineages were measured, including *tumor protein 63 (Trp63), cytokeratin 5 (Krt5), cytokeratin 8 (Krt8),* and *probasin (PB)* (**Fig. 3C and S4**). Marker gene patterns mostly tracked as predicted with either luminal or basal genes based on the GEMM of sample origin. B1 cluster is strongly identified with canonical basal markers. B2 and B3 are fewer in number but also positive for basal markers. In clusters originating from the luminal GEMM, L1 was positive for luminal markers while also strongly expressing basal markers. L2 tracked exclusively to luminal gene patterns and is characteristically PB-expressing. L3, L5, and L6 are relatively small clusters expressing luminal marker transcripts (**Fig. 3C**). L4 differed the most from the other clusters with minimal expression of basal or luminal markers. High in these cells were genes associated with stroma fibroblasts, specifically *Col1a2, Col1a1, Ccl2,* and *Ccl7* (**Fig. 3C and S4**).

We have observed a consistent association of strong PI5P4Kα protein expression with the basal epithelial layer and asked what differences could explain this by comparing the B1 and L2 scRNA-seq clusters. Transcript expression of the three *Pip4k2* isoforms was evaluated (**fig. S5A**). *Pip4k2c* (gamma isoform) transcript is abundant with quantifiable detection across all clusters. *Pip4k2a* and *Pip4k2b* transcripts were in low abundance using the modest sequencing coverage of our data (84, 429 average reads per cell). However, more detectably positive *Pip4k2a-*expressing cells were concentrated in the B1 and L4 clusters compared to the PB-positive L2 clusters (**fig. S5A**). A marker of lysosomal content, Lysosome-associated membrane protein 2 (*Lamp2*), was also upregulated in B1 versus L2 groups (**fig. S5B**). Previously, we and others have associated PI5P4Kα function with lysosomal signaling (*10*, *13*).

To attempt to annotate the clusters, we considered classifications from Crowley *et al*. establishing a scRNA-seq cell atlas of the mouse prostate (*2*). Our single-cell data was collected using a platform different from the 10x genomics approach used by Crowley *et al*., and as expected, merging the two datasets revealed a moderate batch effect (**fig. S5C**). Gene module scores were calculated using the marker genes described in Crowley *et al.* (**table S3**) (*2*). This confirmed B1 as elevated in the basal cell score and low in luminal patterns. Similarly, B3 aligned with patterns of periurethral (PrU) cells, L2 was most similar to AP luminal (LumA), L4 to stroma, L5 to luminal proximal (LumP), and L6 to VP luminal (LumV) (**Fig. 3D**). With less confidence, L3 is likely associated to a dorsal luminal (LumD) cells; however, this group expresses markers of both LumA and LumD. Unexpectedly, clusters B2 and L1 expressed both luminal and basal markers, and neither were strongly associated with annotated clusters from the atlas dataset (**Fig. 3, D and E; S5**). The presence of minor populations of intermediate cells that co-express epithelial markers (i.e., *Krt5+/Krt8+*) have previously been reported from IHC-based cell lineage studies (*28*). Therefore, we classified B2 as ‘basal-intermediate’ and L2 as ‘luminal-intermediate’ clusters. **Fig. 3E** summarizes the annotations assumed to associate with each cluster identified from the two GEMMs.

Overall, these results characterize the cell populations targeted in commonly utilized GEMMs that utilize lineage markers for Cre-based gene deletion (**Fig. 3A**). Three cell clusters were identified from the basal model (CK5-eYFP) with the majority of cells annotated as canonical prostate basal cells (B1) (**Fig. 3B**). These data are valuable for informing the development of a new basal-specific GEMMs, for example to target PI5P4Kα. This instills a level of confidence that the large majority of CK5 positive cells, that experience Cre-mediated gene modifications in the CK5-eYFP model, have uniform basal cell attributes.

### 4. Deletion of Pip4k2a has minor morphological impact on epithelial tissues

PI5P4Kα expression has an inverse relationship with the AR pathway and becomes enriched in the basal layer of prostate glands following castration-induced regression (*10*). We hypothesized that PI5P4Kα signaling in basal cells could support CRPC survival programs. To directly test the impact of downregulating PI5P4Kα in basal cell populations, we crossed CK5-eYFP animals with a GEMM that is homozygote for floxed *Pip4k2a* alleles, referred to as CK5-Pip (**Fig. 4A**). PI5P4Kα was confirmed to colocalize with the CK5 basal cell marker using IF- IHC. Notable, a maximum of ∼30% of CK5+ basal cells were also PI5P4Kα expressors (**fig. S2**). Animals with eYFP+ basal cells with *Pip4k2a^fl/fl^*genotype were compared to CK5-eYFP controls, expressing eYFP but maintaining wild-type *Pip4k2a*. Multiple time points were observed up to 12 months following Cre activation (**Fig. 4, A and B**). It is important to note that the Cre recombinase system is activated in all cells expressing CK5, expressed in several tissues, and not exclusive to the prostate. Tissues sampled from skin, seminal vesicles, testis, bladder, kidney, lung, spleen, pancreas, and liver were evaluated. There were no notable histopathological changes in CK5-Pip in these organs. Differences in prostate gland cellularity were quantified based on the number of nuclei per lobular area. While a trend suggests a potential increase in CK5-Pip prostates, this was not statistically significant (**Fig. 3C**). A single timepoint of animals with a complete genomic loss of the *Pip4k2b* isoform were also examined. Interestingly, the lobes of these prostates appeared cystically dilated when examined histologically and had only 56% of the nuclear density compared to controls (**fig. S6A**). Animals lacking *Pip4k2b* have reduced body weight compared to controls, while additional loss of *Pip4k2a* in CK5+ cells did not enhance these effects (**fig. S6, C and D**).

**Fig. 4.**
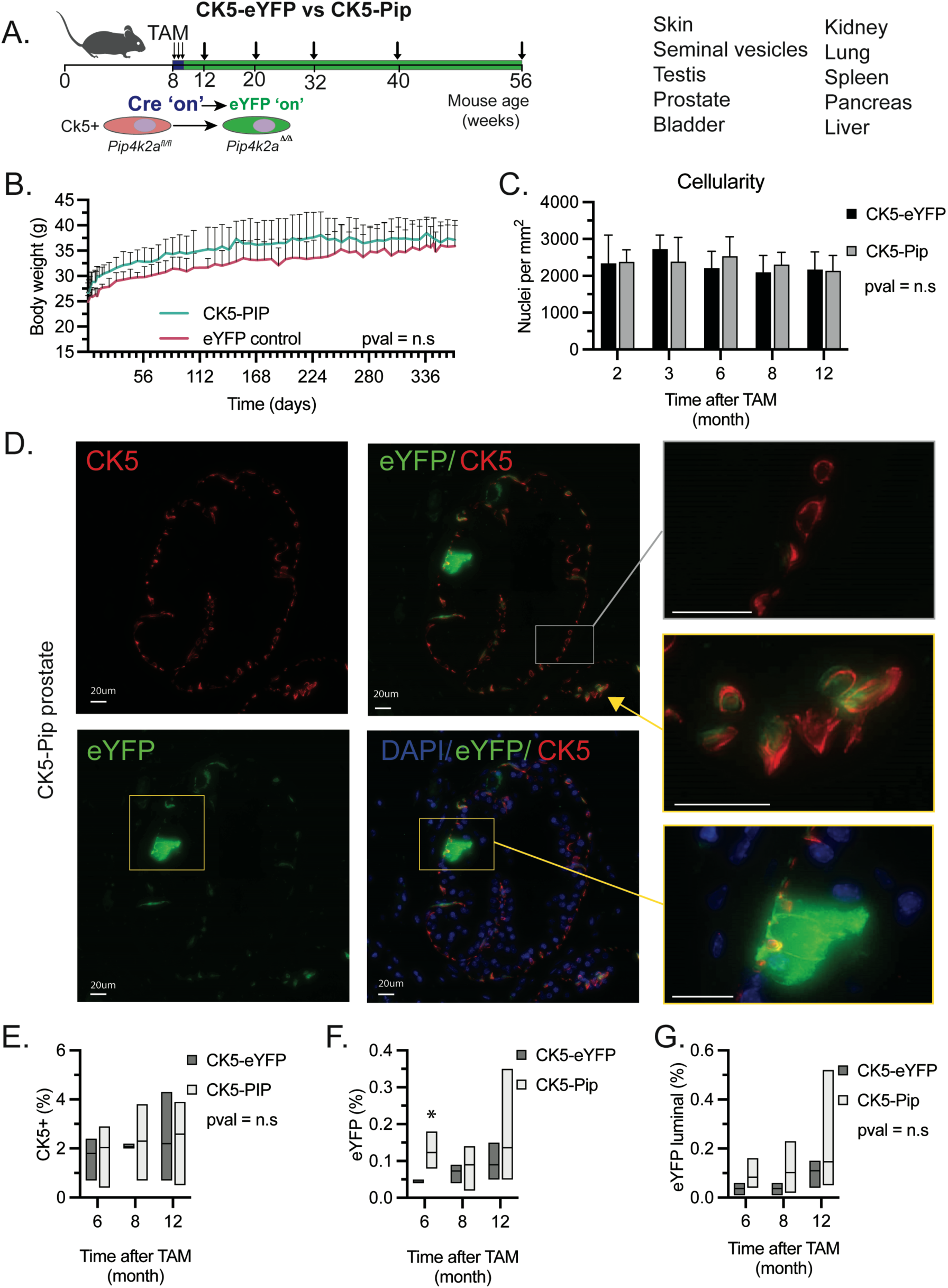
GEMM targeting *Pip4k2a* in CK5+ basal population. (**A**) Graphical summary of *Krt5-Cre^ERT2^; R26eYFP; Pip4k2a^fl/fl^* (CK5-Pip) genetically modified mouse model (GEMM). A homozygote genetic deletion of *Pip4k2a* and expression of fluorescent eYFP protein occurs where Cre is activated. Multiple time points and tissues were evaluated up to 1-year timepoint. No significant differences were measured in (**B**) animal body weight or (**C**) prostate lobe cellularity (anterior lobe represented) compared to CK5-eYFP controls. (**D**) Representative immunofluorescent IHC detection of cytokeratin 5 (CK5) (red) and eYFP Cre (green) reporter with Dapi (blue) nuclear stain in 8-month prostate of CK5-Pip. Zoom regions represent quantified examples of CK5+ basal cells without (grey box) and with (yellow arrow) eYFP expression. Additionally, eYFP+ luminal cell morphology (lower right, yellow box) (scale bar = 20μm). Multi timepoint cohorts of CK5-eYFP and CK5-Pip prostate IF-IHC stain quantification for (**E**) CK5+ cells (pval = n.s), (**F**) percent eYFP positive cells (6-month, 2.85 fold increase, t.test, pval = 0.05), and (**G**) eYFP positive with luminal morphology (pval = n.s). *t* test values: n.s., not significant (*p*> 0.05), * *p* <0.05, ** *p* <0.01, *** *p* < 0.001.

Detection of the Cre reporter, eYFP, was used to consider changes to basal cell lineage in CK5-Pip prostate tissues. The majority of eYFP colocalized with the CK5 basal cell marker with the occurrence of minor populations of eYFP+ cells with a luminal cell morphology (**Fig. 4D**). Digital signaling quantification showed there was no difference in the proportion of basal cells between CK5-Pip and CK5-eYFP controls (**Fig. 4E**). Tissue from 3 individual animals were compared for timepoints 6 and 8, while 5 animals were available for staining and quantification for 12 months timepoint. At six-months CK5-Pip prostate tissue sections had a slight increase in eYFP+ marked cells that was not retained in later timepoints (**Fig. 4F**). Although occurring rarely, the percentage of eYFP+ luminal cells increased on average over time with a slight trend to be higher in the CK5-Pip; however, is not statistically different (**Fig. 4G**). Characterization of CK5-Pip suggests targeting loss of *Pip4k2a* is alone not sufficient to establish oncogenic transformation in basal cells.

### 5. Loss of Pip4k2a slows Pten mutant tumorigenesis

Emerling et al. (2013) demonstrates that loss of PI5P4K isoforms may reverse oncogenesis related to loss of the tumor suppressor *TP53*. Equally as aggressive, the loss of *PTEN* is a major driver of PCa. Therefore, we questioned whether modulating PI5P4Kα impacts disease progression of mPIN driven by *Pten* loss. Previous work has established loss of *Pten* in this CK5+ model produces aggressive adenocarcinoma in the prostate, as well as other tissues (*4*). We generated a double deletion model using *Pip4k2a^fl^* and *Pten^fl^* floxed alleles (referred to as CK5-PipPten). CK5-PipPten were compared to animals with only *Pten* (referred to as CK5-Pten) and observed up to a maximum of three-month timepoint following induction (or 20 weeks of age total) (**Fig. 5A**). In both models, observation of hardened lobe tissue was notable between in two to three month samples following Cre induction, as was increased urogenital mass that is indicative of mPIN (**Fig. 5B and S7A**).

**Fig. 5.**
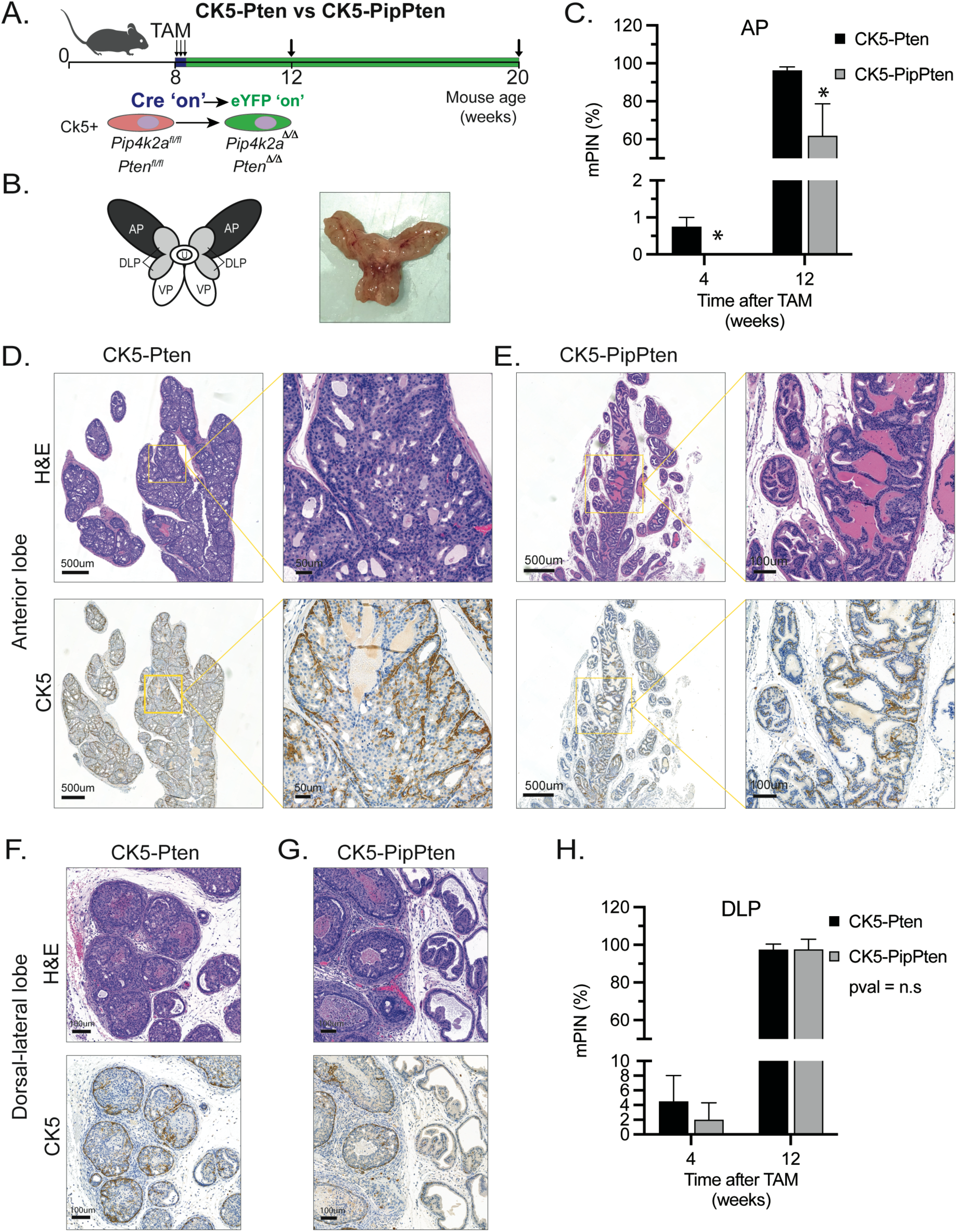
Loss of *Pip4k2a* slows *Pten* mutant tumorigenesis. (**A**) Graphical summary of *Krt5-Cre^ERT2^; R26eYFP; Pten^fl/fl^* (CK5-Pten) and *Krt5-Cre^ERT2^; R26eYFP; Pten^fl/fl^; Pip4k2a^fl/fl^* (CK5-PipPten) genetically modified mouse models (GEMMs). A homozygote genetic deletion of floxed genes and expression of fluorescent eYFP protein occurs where Cre is activated. Timepoints and tissues were evaluated up to 3-month timepoint. (**B**) Representative images of mouse prostate lobes and CK5-Pten mouse prostate sampled with lobes containing dense tumor tissue. (**C**) Anterior prostate (AP) lobes were quantified for the area of mouse prostatic intraepithelial neoplasia (mPIN) at 4 weeks and 12 weeks following Cre activation. CK5-PtenPip cases have reduced mPIN annotated tissue compared to CK5-Pten (4 weeks, −1 fold, pval = 0.02; 12 weeks 0.35 fold, pval = 0.01). Representative hematoxylin and eosin (H&E) and cytokeratin 5 (CK5) IHC images of (**D**) CK5-Pten (scale bar = 20μm and 50μm) and (**E**) CK5-PipPten (scale bar = 20μm and 100μm) AP lobes. Representative H&E and CK5 IHC images of (**F**) CK5-Pten dorsal-lateral (DLP) lobe and (**G**) CK5-PipPten DLP (left) alongside tumor-free ventral lobes (VP) (right). (**H**) DLP lobes were quantified for the area of mPIN at 4 weeks (0.67 fold, pval = 0.67) and 12 weeks (1.0 fold, pval = 0.97) following Cre activation. *t* test values: n.s., not significant (*p*> 0.05), * *p* <0.05, ** *p* <0.01, *** *p* < 0.001.

Prostate tissue collected from 20 mice (CK5-Pten vs. CK5-PipPten) and different time points (1 vs. 3 months), stained with hematoxylin and eosin (H&E), AR, CK5, and PI5P4Kα, was examined for any relevant histopathologic features. The most evident histologic feature consisted of extensive mPIN in the 3-month groups and variably distended (predominantly ventrolateral) glands in animals corresponding to the one-month time point. The extent of mPIN varied between the different prostate regions. In the anterior gland, CK5-Pten mice presented with more extensive mPIN than the CK5-PipPten group (**Fig. 5, C to E**). This observation could prompt speculation of an oncogene role of *Pip4k2a*. The situation differed for the dorsal-lateral (DLP) glands, which presented with widespread mPIN, without evident differences between CK5-PipPten and CK5-Pten mice (**Fig. 5, F to H**). **Fig. 5G** demonstrates a side-by-side comparison of DLP (left) and VP (right) lobes that represent both models. While DLP is consumed with advanced mPIN, the VP retains a high proportion of benign glandular tissue. Given the only rare and mild effect on the ventral and dorsal glands, the development of mPIN appears to have a clear preference for the lateral and anterior lobes in this model. At one month, most mice presented with small mPIN foci or precursor lesions, primarily seen in the lateral gland, confirming the lateral location of initial mPIN development (**Fig. 5, C and H**). Evidence for invasive neoplastic changes were characterized by rare foci of mPIN microinvasion. Even though only ill-defined, this feature tended to be less frequent in CK5-PipPten mice, supporting the role of PI5P4Kα in promoting malignant transformation. Comedonecrosis and a variable level of stromal inflammation were common in mPIN-affected prostate regions. These features were most pronounced in the lateral gland and were more evident with increasing extent of mPIN. Loss of PTEN protein was confirmed in all lobes of CK5-PipPten prostates, as was the progression of basal cell hyperplasia to neoplastic luminal differentiation, which was originally characterized in the CK5-Pten GEMM (**fig. S7, B and C**). Altogether, the loss of *Pip4k2a* in basal cells suppresses the early stages of mPIN progression of aggressive homozygous *Pten* mutant PCa.

### 6. In vivo loss of Pip4k2a triggers shift in metabolic pathways

Having optimized our approach for targeted-selection scRNA-Seq from *in vivo* models (**Fig. 3**), we next asked what changes occur in basal cells following the loss of *Pip4k2a.* Three 96-well plates were collected from two CK5-eYFP control animals and two CK5-Pip animals after Cre induction using eYFP+ FACs at a two-month timepoint. Following quality control, pathway changes between control and *Pip4k2a* loss were analyzed for 2210 and 1478 cells, respectively (**Fig. 6A, S8A; table S4**). Strong downregulation was observed in metabolic pathways associated with functional studies on PI5P4Ks, including a decrease in oxidative phosphorylation (NES −2.23, padj = 4.8E-11), fatty acid metabolism (NES −1.98, padj = 1.4E-05), glycolysis (NES −1.55, padj = 6.3E-03), and cholesterol homeostasis (NES −1.58, padj = 2.4E-02). Notable upregulated pathways include MYC targets V1(NES 2.55, padj = 3.3E-17), P53 pathway (NES 2.15, padj = 2.8E-09), and the unfolded protein response (NES 1.85, padj = 1.5E-04). Transcript expression from *in vivo* models shows a clear pattern of downregulated metabolism pathways and enhanced activation of known cell stress compensatory pathways (i.e., MYC) in CK5-Pip basal cells.

**Fig. 6.**
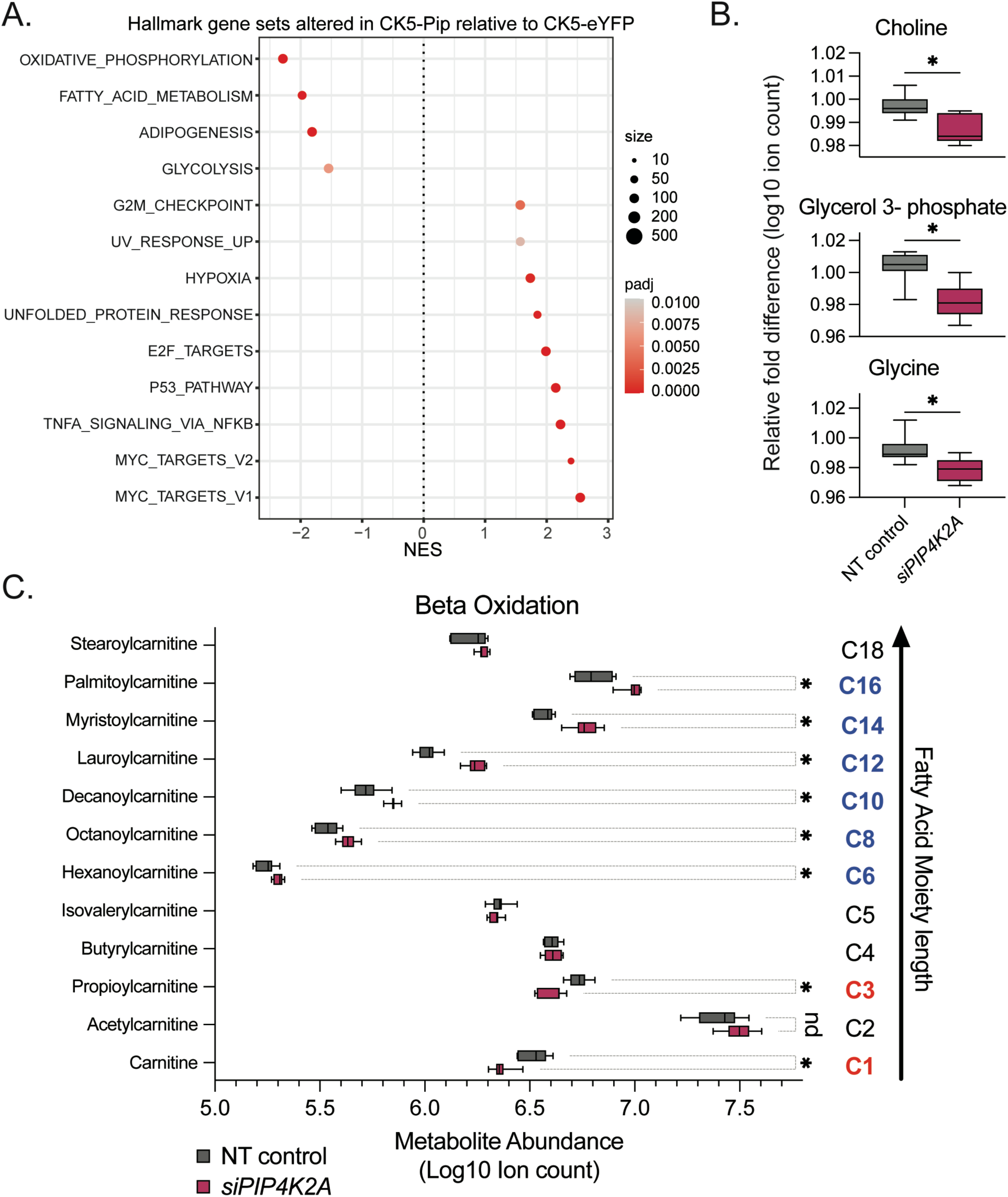
Targeting *Pip4k2a* triggers a shift in metabolic pathways. The differences in transcript expression of CK5-Pip eYFP+ prostate cells were compared to CK5-eYFP controls sampled *in vivo* at a 2-month timepoint using SORT-seq scRNA sequencing. (**A**) Gene set expression analysis (GSEA) of most altered Hallmark pathways are shown with pval <0.01 cut-off. The LNCaP human PCa cell line was treated *in vitro* with siRNA targeting *PIP4K2A* or a non-targeting (NT) control in conditions of hormone-stripped medium and evaluated using targeted metabolomics. (**B**) Key metabolites linked to lipid regulation are highlighted, and the extended list is shown in **fig. S8 and table S5**. (**C**) Most noteworthy were changes to carnitine metabolites associated with lipid beta-oxidation. Enrichment of carnitines with lipid moieties in medium and long (C6-C18) acyl chain length observed following *siPIP4K2A* knockdown and depletion of C1 and C3 short acylcarnitines. *t* test values: n.s., not significant (*p*> 0.05), * *p* <0.05, ** *p* <0.01, *** *p* < 0.001.

Finally, to validate our scRNA-Seq observations on a human PCa model, targeted metabolomics was performed on LNCaP cells (PTEN mutant) following siRNA knockdown of *PIP4K2A* in conditions of androgen depletion. Knockdown conditions were compared to a non-targeting (NT) RNA control and an untreated medium control (**fig. S8, B and C, table S5**). *PIP4K2A* depletion resulted in a decrease in glycerophospholipid metabolites, choline (log2 −0.53, pval = 2.3E-02), glycerol 3-phosphate (log2 −0.37, pval = 1.0E-02), and glycine (log2 −0.26, pval = 1.5E-02) (**Fig. 6B**). Importantly, another indication of dysfunctional lipid regulation was detected in the analysis of acyl chain linked carnitine species (**Fig. 6C**). Twelve carnitines with fatty acid moiety lengths ranging from C1-C18 were compared between *siPIP4K2A* and NT control. Two short-chain lipids (C1 and C3) were decreased. However, six of seven medium to long-chain carnitines (C6-C18) were significantly enriched). PCa metabolism is uniquely fueled by the oxidation of lipid species, which are transported between organelles using carnitine shuttle systems (*29*, *30*). The enrichment of carnitine metabolites supports a block in fatty acid metabolism pathways following loss of PI5P4Kα.

## DISCUSSION

The efficacy of precision medicine-guided treatments depends on the molecular context for which a therapy is best suited. Here, we consider how cell identity influences the phenotype related to the downregulation of PI5P4Kα, a PI kinase presently considered a cancer drug target. We show that (i) PI5P4Kα has strong but sparse expression in basal cells, and (ii) *in vivo* loss of *Pip4k2a* slows mPIN in combination with PTEN loss. Further, (iii) we suggest this could result from downregulated lipid metabolism pathways and hypothesize loss of *Pip4k2a* impairs the trafficking of lipids needed for beta-oxidation, a process known to be orchestrated by PI5P4Kα (*10*, *13*, *15*). These observations build upon growing evidence that nominates PI5P4Kα as a therapeutic target for PCa. Expanding preclinical efforts are underway to generate PI5P4K inhibitors, which offer new opportunities for oncology drug development (*18*).

We found a loss of PI5P4Kα slowed PCa phenotypes, consistent with reports in breast cancer, AML, and sarcoma (*15*, *19*, *31*). Notably, these results are contrary to observations made in GBM. Shin *et al.* (2019) identified PI5P4Kα in an unbiased screen as having tumor suppressor attributes and proposed that PI5P4Kα competitively binds with p85 to regulate the PI3K signaling cascade. While we do not see this p85 binding relationship in our models (data not shown), multiple networks may be impacted by PI5P4Kα dysregulation, such as direct regulation of mTORC1/2, autophagy, and lipid trafficking (*13*, *32*). The exact role of PI5P4Kα may be dictated by tissue attributes. While cells in both the prostate and the brain rely on lipid metabolism network, these processes are linked to specialized functions. For example, prostate luminal cells upregulate fatty acid oxidation as an energy source because the citric acid cycle (TCA) is rewired to produce and secrete citrate (*33*). In the brain, neurons and glial cells have a lower rate of fatty acid oxidation and higher reliance on glycolysis, a great deal of complex lipid synthesis and storage from astrocytes is committed to supporting neuronal myelination (*34*). Notably, transcript expression of *PIP4K2A* is elevated in neural progenitor cell populations (*20*). Epithelial progenitor cell populations are enriched in the basal layer of prostate glands, where we observed PI5P4Kα positivity, thereby opening a potential line of investigation. In future work, there could be great value in examining the role of PI5P4Kα, specifically in stem cell or progenitor populations.

The genetic deletion of *Pip4k2a* alone was not sufficient to dramatically modify prostate gland morphology. However, a phenotype was observed in combination with PTEN tumor suppressor mutation. A combination of altered PTEN is a common approach in PCa GEMMs (*35*, *36*). Relevant to our findings, the deletion of the fatty acid translocate, *Cd36,* in a *PB-Cre; Pten^fl/fl^*mouse model demonstrated that PCa develops slower following inhibition to fatty acid uptake (*37*). CD36 is frequently upregulated in clinical samples and exemplifies the importance of lipid trafficking PCa. Although the CK5-PipPten model has a similar delay in *Pten* mutant cancer progression, *PIP4K2A* genomic alterations are considerably rare in clinical cases. However, it cannot be ruled out that PI5P4Kα could be activated at other regulatory steps to benefit oncogenesis. It is important to note that neither the expression nor genomic alteration of *PIP4K2A* correlates with *PTEN* mutation in human samples (*10*). Based on these assumptions, the influence of PI5P4Kα on cancer-enabling metabolism may be overlooked in large consortium analyses of multi-omic clinical data.

In this work, we observed diverse PI5P4Kα protein expression patterns across luminal and basal prostate cells. These variations in tissue made us question if important biology is overlooked when characterizing PI5P4Kα in only *in vitro* models. Based on this, we adopted scRNA-seq methodology with two major goals. First, to analyze how *Pip4k2a* loss impacts a specific cell type sorted directly from tissue. Only one-third of basal cells appeared enriched with PI5P4Kα using IF-IHC, making a single-cell resolution particularly necessary. Unfortunately, due to the low-depth coverage of current scRNA-Seq technologies, future experiments will require higher reads coverage to characterize *PIP4K2A* transcripts comprehensively. Our second goal was to characterize the true cell populations impacted by our GEMM systems. Up to this point, there have been general assumptions that PB-driven Cre systems modify mostly luminal populations as well. Cre activation of CK5+ populations is assumed to target basal populations. Our analysis mostly paralleled populations previously defined in scRNA-Seq atlas studies (*2*). Whereas the mouse luminal cells clustered in groups with lobe-specific transcript signatures, the basal cell populations were primarily a single uniform group. In future work, it would be valuable to characterize differences between the CK5-Pten and Ck5-PipPten models using scRNA-Seq and metabolomic technologies.

This study demonstrates the importance of considering cell type when exploring metabolic regulators like PI5P4Kα. The molecular profiles of PCa are frequently overlayed with attributes of luminal or basal epithelial cells to predict disease progression (*9*). A growing body of evidence links epithelial lineages with distinct patterns of metabolism. For example, modulating the rate of pyruvate oxidization and lactate accumulation is sufficient to drive luminal cell differentiation from basal progenitor cells. These resulting metabolite shifts are shown to initiate lineage-defining chromatin remodeling (*38*). Comparably, our models targeting PI5P4Kα also shifts nutrient pathways. While there was no difference in the percentage of lineage-traced luminal cells from the CK5-Pip model, the metabolic shift resulting from *Pip4k2a* loss may influence the basal to luminal differentiation that occurs in basal cell origin PCa (*6*). A blockade in lineage transition might contribute to the deceleration in cancer phenotypes.

Evidence suggests that loss of PI5P4Kα leads to disruptions in fatty acid metabolism and stall in long fatty acid beta-oxidation. Lipid synthesis and oxidation processes are heavily investigated for PCa therapeutics due to their upregulation during progression. For example, inhibition of fatty acid synthase (FASN), a multienzyme protein responsible for catalyzing fatty acids like palmitate (C16:0), is being evaluated in clinical trials for CRPC (NCT04337580) (*39*). In addition, dietary supplementation with sulforaphane (SFN) is being tested for chemopreventative properties, also in PCa (NCT03665922). Preclinical experiments suggest SFN can inhibit FASN and carnitine palmitoyltransferase 1 (CPT1A), enabling fatty acid uptake to the mitochondria for beta-oxidation (*40*). The cancer-delaying phenotype of PI5P4Kα inhibition could parallel the promise of FASN inhibitors or SFN. Specifically, loss of PI5P4Kα *in vivo* suppressed fatty acid metabolism pathways and slowed the progression of the especially lipogenic PTEN mutant PCa GEMM.

Tool compounds that inhibit PI5P4K isoforms are currently being explored (*16*, *18*). It is intriguing to consider a PI5P4Kα inhibitor for PCa chemoprevention. *Pip4k2a* genomic knock-out mice are essentially without phenotype, suggesting tolerability. Most men with localized prostate cancer are diagnosed in the seventh decade of life and usually die of other causes. Therefore, chemoprevention has always been an attractive strategy to slow down cancer progression in an elderly patient in combination with active surveillance. Finasteride demonstrated promise but was ultimately not adopted (*41*). Given the common loss of PTEN in early prostate cancer, the use of a PI5P4Kα inhibitor might have a role in a chemopreventive strategy where the goal would be to delay tumor growth.

In conclusion, PI5P4Kα is enriched in prostate basal cells, a hub of regenerative progenitor cells, and a potential cell of origin of PCa. Understanding the molecular underpinnings of the basal layer offers insights into cellular programs hijacked in malignancy. This work exemplifies the importance of capturing rare cell signaling biology from intact *in vivo* systems that may not be fully reflected using cell culture systems. Finally, we suggest targeting PI5P4Kα in early neoplastic development could be a means to slow *PTEN* mutant PCa.

## MATERIALS AND METHODS

### Human prostate dataset queries

The ‘*A Cellular Anatomy of the Normal Adult Human Prostate and Prostatic Urethra*’ dataset was analyzed from publicly available data from the 10x Genomics 3’ V2 platform. 28 702 human prostate cells were subset for cells with positive detection of *PIP4K2A* transcript with cell type annotations as previously described (accessible at https://cellxgene.cziscience.com/e/c3fe3c1e-5bf8-4678-b74a-79899243ad41.cxg/) (*25*, *42*).

### Animal care and housing

All animal studies were approved by the Cantonal Veterinary Ethical Committee, Switzerland (license BE46/2021). Animals were housed in ventilated cages with unrestricted access to pre-sterilized food and fresh water. A maximum of 5 animals were maintained per cage on Aspen bedding. Ambient temperature was 20±2 °C, kept at a constant humidity of 50±10% and on a 12-hour automatic light-dark cycle.

### Genetically engineered mouse models (GEMM)

*Pip4k2a^tm1.2Lca^* animals (MGI:5568930), referred to as *Pip4k2a^flox^ (flox* referred to as *fl)*, were a kind gift of the group of Lewis C. Cantley. The generation of *Pip4k2a^fl^* (*13*, *19*), *CK5-CreER^T2^* (*43*), *Probasin-Cre* (PB-Cre) (*44*), *Pten^fl^* (*45*), and *Rosa26eYFP* (*46*) mice have been previously described. These strains were then crossbred to generate the CK5-eYFP (*Krt5-CreER^T2^; R26R-eYFP*), CK5-Pip (*Krt5-CreER^T2^; R26R-eYFP; Pip4k2a^fl/fl^*), CK5- Pten (*Krt5-CreER^T2^; R26R-eYFP; Pten^fl/fl^*), CK5-PipPten (*Krt5-CreER^T2^; R26R-eYFP; Pip4k2a^fl/fl^; Pten^fl/fl^*), and PB-Cre4; R26eYFP mice used in the mouse study. Animals were genotyped using the Transnetyx real-time PCR platform and standard PCR assays. For *CK5-CreERT2* Cre recombinase-inducible models, animals were gavaged for three consecutive days with 75 mg/kg tamoxifen citrate prepared in filter-sterilized corn oil at eight weeks of age (total dose 225mg/kg). A summary of genetically modified animal lines is found in **table S1**.

### Immunohistochemical protein detection

Tissue sections were processed using standard histology procedures, fixed with 4% formaldehyde overnight, and paraffin-embedded, and sections were cut at 2.5 μm. Conditions for additional antibodies are listed in **table S2**. As previously described, slide staining was done using the BOND RX autostainer (Leica Biosystems) (*10*).

### Immunofluorescent staining

Adult C57BL/6JRj mouse prostates were harvested and paraffin-embedded. Tissue serial sections were deparaffinized in xylene and rehydrated using decreasing ethanol concentrations (100 to 95 to 70% diluted in H_2_O). Heat-induced epitope retrieval was used at 98°C in tris-EDTA buffer. Slides were left to return to room temperature, washed in Tris-buffered saline (TBS), blocked in 5% normal goat serum at room temperature, and probed with primary antibodies in an immunostain moisture chamber. Protein targets were detected using optimized conditions (**table S2**). DAPI (100 ng/ml) was used as a nuclear counterstain, and mounting was done with ProLong Gold (Thermo Fisher Scientific; P36930). Images were acquired with 40x oil immersion on a Zeiss LSM 710 confocal microscope and/or with 20x magnification on a Panoramic 250 (3DHistech) slide scanner.

### Digital tissue quantification and pathological evaluation

Whole-slide images were acquired with a Panoramic 250 (3DHistech) slide scanner and exported from CaseView software (version 2.5). Coded file names were used to blind the investigator from the experimental group during data acquisition and annotation. Also, sample group blinding was implemented for the evaluation of tissue samples by pathologists. Multiple tissues, including a priority on prostate tissue, from each mouse was stained for HE, CK5, and Pip4k2a. Digital histopathologic evaluation was performed by a board-certified veterinary pathologist (Dipl. ECVP). Tissue slides were evaluated for any histomorphologic change. On HE slides, the presence of distended glands and mouse prostate intraepithelial neoplasia (mPIN) was quantified using Visiopharm (Visiopharm, Horsholm, Denmark) software with the following workflow: 1) Manual annotation of the region of interest (ROI), defined as prostate tissue; 2) automated detection of gland profiles (deep learning-based classification); 3) manual assignment of altered (distended or mPIN) vs. non-altered gland profiles and corresponding lobe (AP, DLP or VP); 4) Output: area of altered vs. non-altered glandular tissue, separately for the different prostate regions [absolute (mm^2^) and relative (%)]. For each case, the prostate was semi-quantitatively assessed for mPIN microinvasion, mPIN comedonecrosis, and stromal inflammation, using the following scoring: 0: absent or neglectable; 1: mild; 2: moderate; 3: severe. The cellularity of the AP was quantified using Visiopharm software with the following workflow: 1) Manual definition of the anterior prostate region as ROI; 2) automated detection of gland profiles (deep learning classification) with manual correction where needed; 3) automated detection and labeling of nuclei (deep learning classification); 4) output: total number of nuclei; area of ROI (mm^2^); cellularity (number of nuclei per mm^2^).

### In vitro 3D organoid cultures

Mouse prostate organoids were generated as previously described (*10*, *47*). Adult anterior lobe prostates were harvested, dissociated, and FACS-sorted for the eYFP^+^ cell population from mouse lines containing activated *R26eYFP* Cre recombinase reporter (**Fig. 2C**). Organoids were maintained in 3D Matrigel culture below ten passages to limit cell identity drift.

### Protein Western Blotting

PBS-washed cell pellets were extracted for protein lysates using radioimmunoprecipitation assay buffer (RIPA) with protease and phosphatase inhibitors. Total protein concentration was measured using the Pierce BCA Protein Assay Kit (Thermo Fisher Scientific). Between 20 and 50 μg of protein was used for standard SDS– polyacrylamide gel electrophoresis procedures with Mini-Protean TGX gels (4 to 15%; Bio-Rad, 456-1084). An iBlot2 quick transfer system (Thermo Fisher Scientific, IB23001) was used to transfer protein to the nitrocellulose membrane. Blocking and secondary antibody incubations were done using 5% milk in Tris-buffered saline with 0.1% Tween^®^ 20 detergent, and primary antibodies were probed with optimized conditions (**table S2**). Three 5-min wash steps were used between incubations and before visualization. Protein band detection was done with chemiluminescence using the Luminata Forte substrate (Thermo Fisher Scientific, WBLUF0100) and acquired with the FUSION FX7 EDGE Imaging System (Witec AG). ImageJ version 1.53a was used to measure band densitometry along with normalization to housekeepers.

### SORT-Seq single-cell RNA sequencing

Whole mouse prostates were harvested and digested to single cell solution as previously described (*10*, *47*) using 5mg/mL collagenase in advanced DMEM (+/+/+) for 1 hour of shaking at 37 degrees Celsius. This was followed by medium washes and 15-minute incubation with TrypLE at 37 degrees Celsius then the digest solution was passed through a 70uM filter. Cells treated with DAPI for live/dead stain were sorted at the Department of BioMedical Research FACs Core facilities in a solution of PBS with 0.2% fetal bovine serum. Protocols for sorting single eYFP+ cells into wells of 384-well SORT-Seq plates were conducted under the recommendations of Single Cell Discoveries (https://www.scdiscoveries.com/). eYFP positive control prostates were used to adjust A488 gating settings for each sort. Two animals and three 96-well plates were collected as single cells for each group, centrifuged at 4 degrees Celsius, snap-frozen on dry ice, and then stored at −80 degrees Celsius until transport to sequencing facilities. The SORT-seq protocol was used to generate libraries for scRNA-Seq, a partially robotized method based on the CEL-Seq2 protocol (*24*, *48*). Standardized protocols for library preparation and sequencing reads alignment has previously been summarized (*49*).

### Single-cell data analysis

Data were analyzed using the Seurat package v.4.0. 6 (*27*). Cell QC filtering was done using the following thresholds: nCount > 500, nCount < 40000, nFeature > 300 percent.mito < 20, log10GenesPerUMI > 0.8 percent.spike_in < 30. Differential gene expression analysis between clusters was done using the FindAllMarkers function from the Seurat package. Differential gene expression between Pip4k2a-loss cells (CK5-Pip) and control cells (CK5-eYFP) was done with DESeq2 v.1.34.0 (*50*) through the R package DElegate v.1.0.0 (*51*) Gene set enrichment analysis was done using the package fgsea v.1.20.0 (*52*) and the mouse gene sets from the Molecular Signatures Database (https://www.gsea-msigdb.org). Analysis was done in R software (v.4.1.2).

Single cell data from Crowley *et al.* (*2*), were downloaded from the GEO repository (accession number GSE150692). All mouse samples were merged and processed with the Surat package as described above, and then merged with our data. Marker genes for each subtype were chosen from Figure 1—figure supplement 4 (*2*)(**table S3**) and were used to calculate a per-cell gene set score using the ModuleScore function from the Seurat package. A small cluster of cells from Crowley *et al.* expressed both LumA and LumD markers, so we labeled it LumAD (not described in the original paper).

### Metabolomics

Metabolite extraction was conducted with the addition of MeOH: H2O (4:1) on samples from four experimental groups (LNCaP RPMI, charcoal-stripped (CS), NT, and siPIP4K2A), each with six biological replicates. Samples were homogenized, centrifuged with the supernatant, and then evaporated with a vacuum concentrator (LabConco, Missouri, US). For protein quantification, pellets were evaporated and lysed in 20 mM Tris-HCl (pH 7.5), 4M guanidine hydrochloride, 150 mM NaCl, 1 mM Na2EDTA, 1 mM EGTA, 1% Triton, 2.5 mM sodium pyrophosphate, 1 mM beta-glycerophosphate, 1 mM Na3VO4, 1 μg/ml leupeptin. Total protein concentration was determined using the BCA Protein Assay kit (Thermo Scientific, #23225). Before LC-MS/MS analysis, dried extracts were resuspended in MeOH: H2O (4:1, v/v) according to protein content.

LC-MS analysis was designed for targeted acquisition of multiple pathways. Samples were analyzed by Hydrophilic Interaction Liquid Chromatography coupled to tandem mass spectrometry (HILIC -MS/MS) in both positive and negative ionization modes using a 6495 triple quadrupole system (QqQ) interfaced with 1290 UHPLC system (Agilent Technologies) (*53*). For quality control, pooled sample aliquots were analyzed periodically throughout the overall analytical run to assess the data quality, correct the signaling intensity drift, and remove peaks of poor reproducibility (CV>30%). Additionally, diluted quality control series were prepared with a methanol solvent and evaluated at the beginning and end of the sample batch. Raw LC-MS/MS data was processed using the Agilent Quantitative analysis software (version B.07.00, MassHunter Agilent technologies). Relative quantification of metabolites was based on EIC (Extracted Ion Chromatogram) areas for the monitored MRM transitions. Signal intensity drift correction was done within the LOWESS/Spline normalization program (*54*) followed by filtering of “not-well behaving” peaks (CV (QC peaks) > 25% & R2 (QC dilution curve) < 0.75) on the reported peak areas of detected metabolites using “R” software R software (v.4.1.2).

### Statistics and data availability

Data are expressed as means ± standard deviation (SD). As indicated, statistical analyses for all data, including microscopy quantification and animal measurements, were performed using students’ two-tailed t-test or ANOVA using GraphPad Prism. Statistical significance is indicated on figures (*p<0.05, **p<0.01, ***p<0.001, n.s., not significant p >0.05) unless specified otherwise. The datasets generated during and/or analyzed during the current study are available from the corresponding author upon reasonable request. Single-cell RNA sequencing data will be deposited publicly upon publication.

## SUPPLEMENTARY INFORMATION

Supplemental figures S1 – S8

Supplemental tables S1 and S2

Supplemental tables S3-S5 (see Excel files)

- Crowley *et al.* markers for scRNA-Seq module scores
- GSEA pathways from Fig. 6
- Metabolomics summary

## ACKNOWLEDGEMENTS

We especially appreciate the kind help from our entire team and collaborators. This includes the University of Lausanne Metabolomics Unit, the University of Bern Microscopy Imaging Center (MIC), and the University of Bern Central Animal Facilities (CAF). We acknowledge indirect support from the Bern Center for Precision Medicine (BCPM) pilot programs. As well the labs of Dr. Lewis C. Cantley and Dr. Marianna Kruithof-de Julio for sharing of animal resources.

## FUNDING

This work was funded by the Swiss National Science Foundation (SNF#31003A_175609, SNF#310030_207635), an EU Commission Marie Sklodowska-Curie Individual Fellowship (PCAPIP), and The Johanna Dürmüller-Bol Foundation.

## AUTHOR CONTRIBUTIONS

Conceptualization: JT, BME, MAR

Methodology: JT, ML, MR, AB, SdB, MAR

Investigation: JT, AB, SdB

Visualization: JT, AB

Funding acquisition: JT, MAR

Project Administration: JT, MAR

Supervision: MAR and SKYN

Writing-original draft: JT, AB, MAR

Writing-review & editing: all authors

## COMPETING INTERESTS

Authors declare that they have no competing interests.

## DATA AVAILABILITY

All other data are available in the main text or the supplementary materials. Data generated by scRNA-Seq will be made available publicly at time of publication.

## Notes

### Competing Interest Statement

The authors have declared no competing interest.

