## Supplemental Figures and Tables S1-S3 for "Loss of PI5P4Kα slows the progression of a *Pten* mutant basal cell model of prostate cancer"

**Supplemental figures S1 – S8**

**Supplemental tables S1 and S2**

**Supplemental tables S3-S5 (see Excel files)**

- **Crowley *et al.* markers for scRNA-Seq module scores**
- **GSEA pathways from Fig. 6**
- **Metabolomics summary**

**Supplemental Figures**

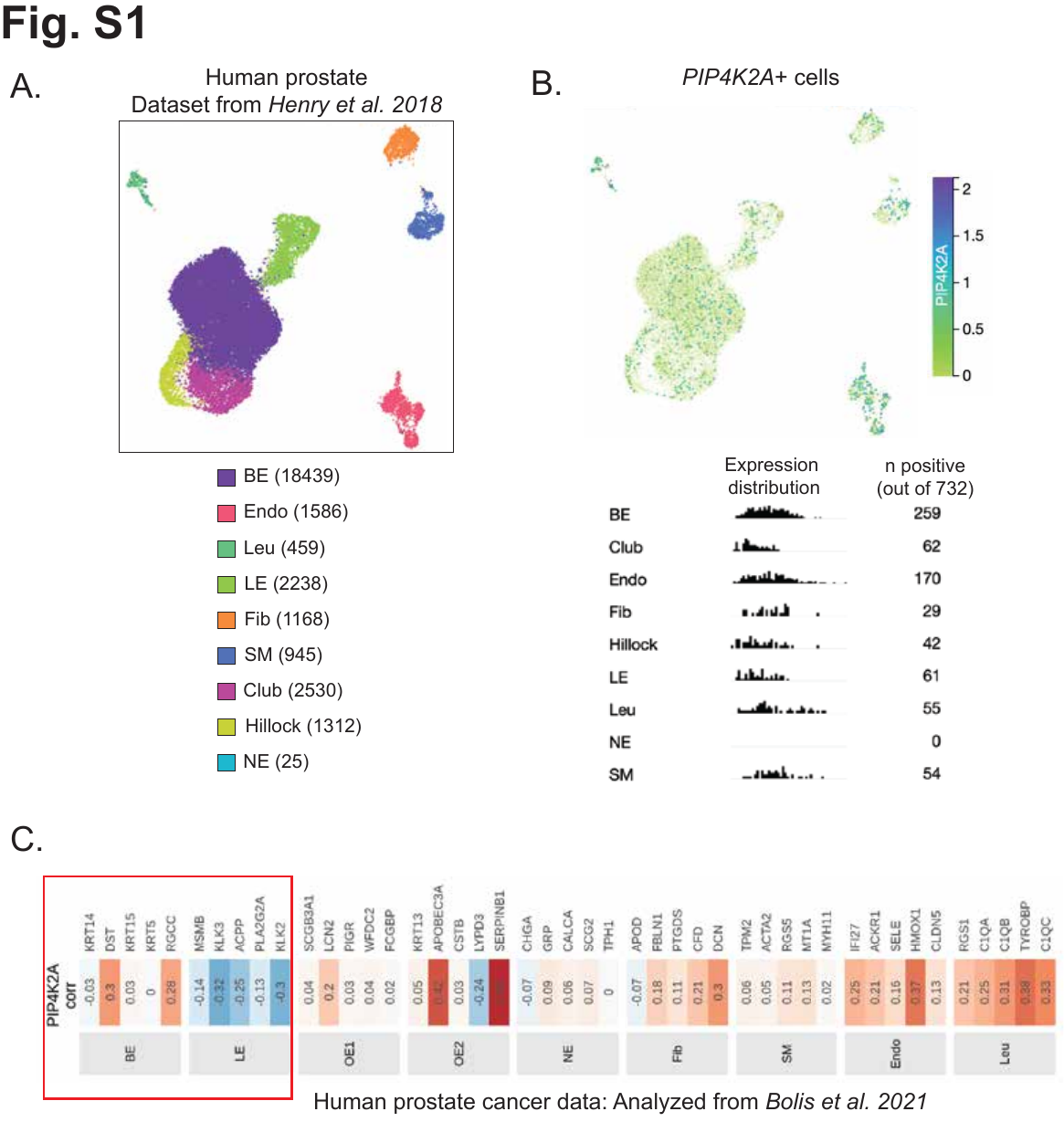

**Fig. S1. *PIP4K2A* expression in human scRNA-Seq dataset.** (**A**) A human adult prostate single cell RNA-Seq (scRNA-Seq) dataset is annotated for basal epithelium (BE), luminal epithelium (LE), club, endothelia (Endo), fibroblasts (Fib), hillock, leukocytes (Leu), smooth muscle (SM), and neuroendocrine (NE) populations (*25*, *41*). Numbers in brackets represent the number of cells for each group. (**B**) Detection of *PIP4K2A* transcript is demonstrated by UMAP and normalized expression distribution plots based on cell type annotation. (**C**) Genes associated with cell type classification correlated to *PIP4K2A* transcript expression in human PCa data set from *Bolis et al., 2021.* Relative differences between BE and LE enrichment highlighted in the red box

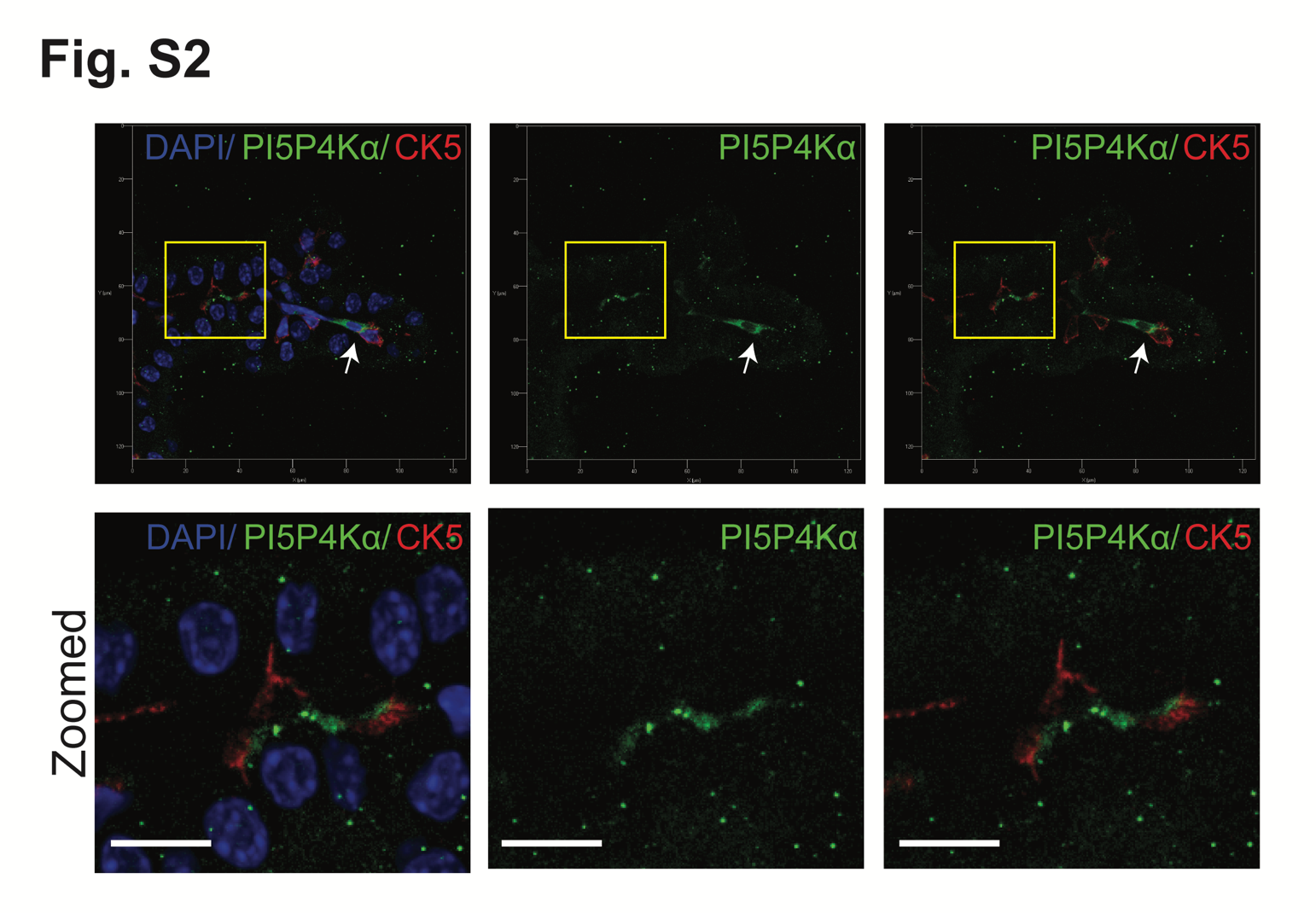

**Fig. S2. PI5P4Kα expression enriched in basal compartment.** Immunofluorescent detection of PI5P4Kα and cytokeratin 5 (CK5) basal cell marker in adult mouse prostate tissue. Signals both localize to basal layer. Zoomed region represented in yellow box. Strong positive PI5P4Kα indicated with white arrow.

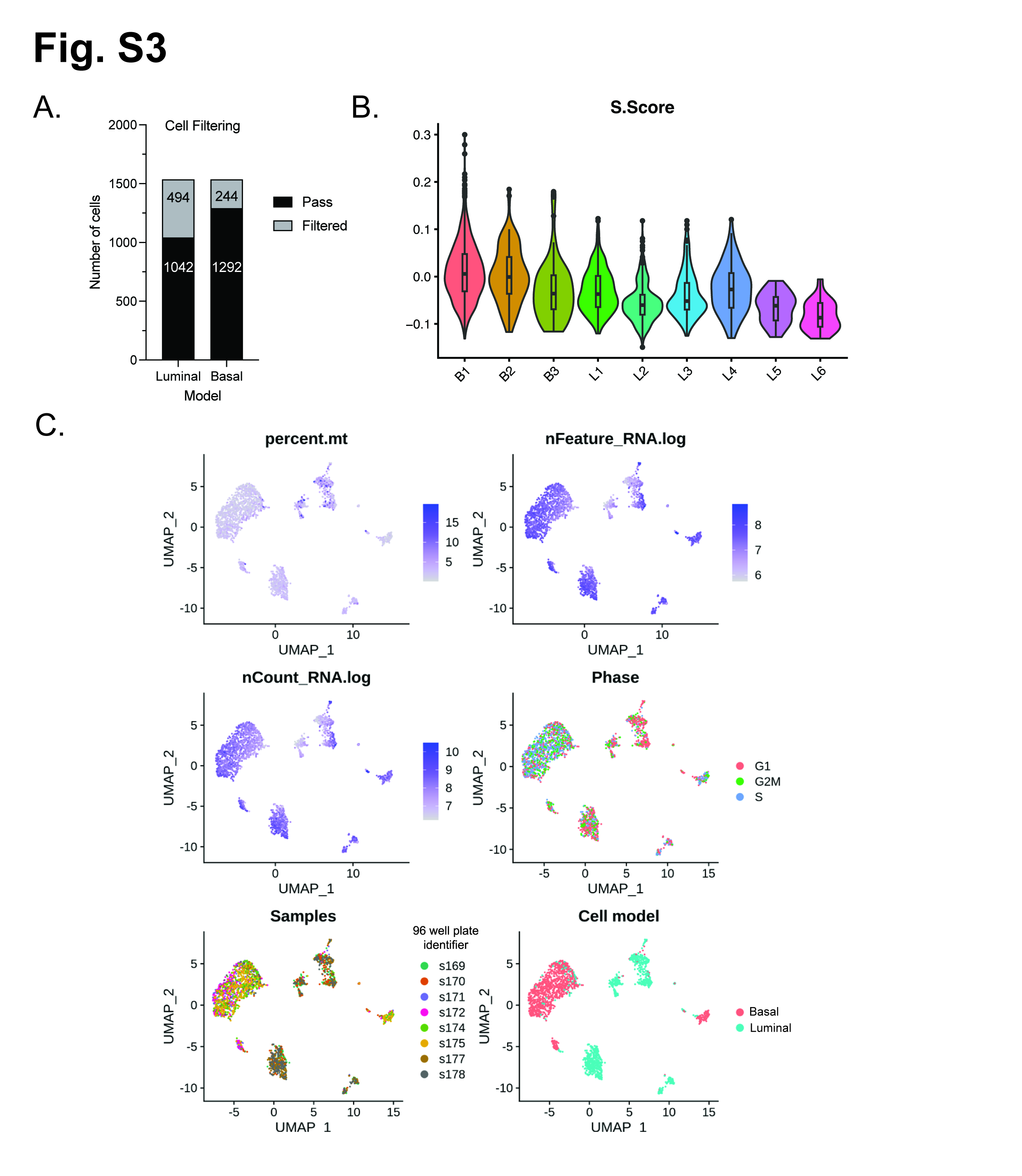

**Fig. S3. Selected cell scRNA-Seq evaluation of prostate GEMMs.** In basal and luminal GEMMs from **Fig. 3** eYFP+ cells were collected with FACs and (**A**) 1500 cells per model were evaluated for data quality, with 69% luminal and 86% basal model cells passing quality filtering. (**B**) Violin blots of cluster distribution of S phase, or cell cycling populations. (**C**) Summary UMAPs of percent mitochondrial genes, RNA content, cell phase, sample distribution (by 96-well plate), and origin of cell based on mouse model (basal or luminal).

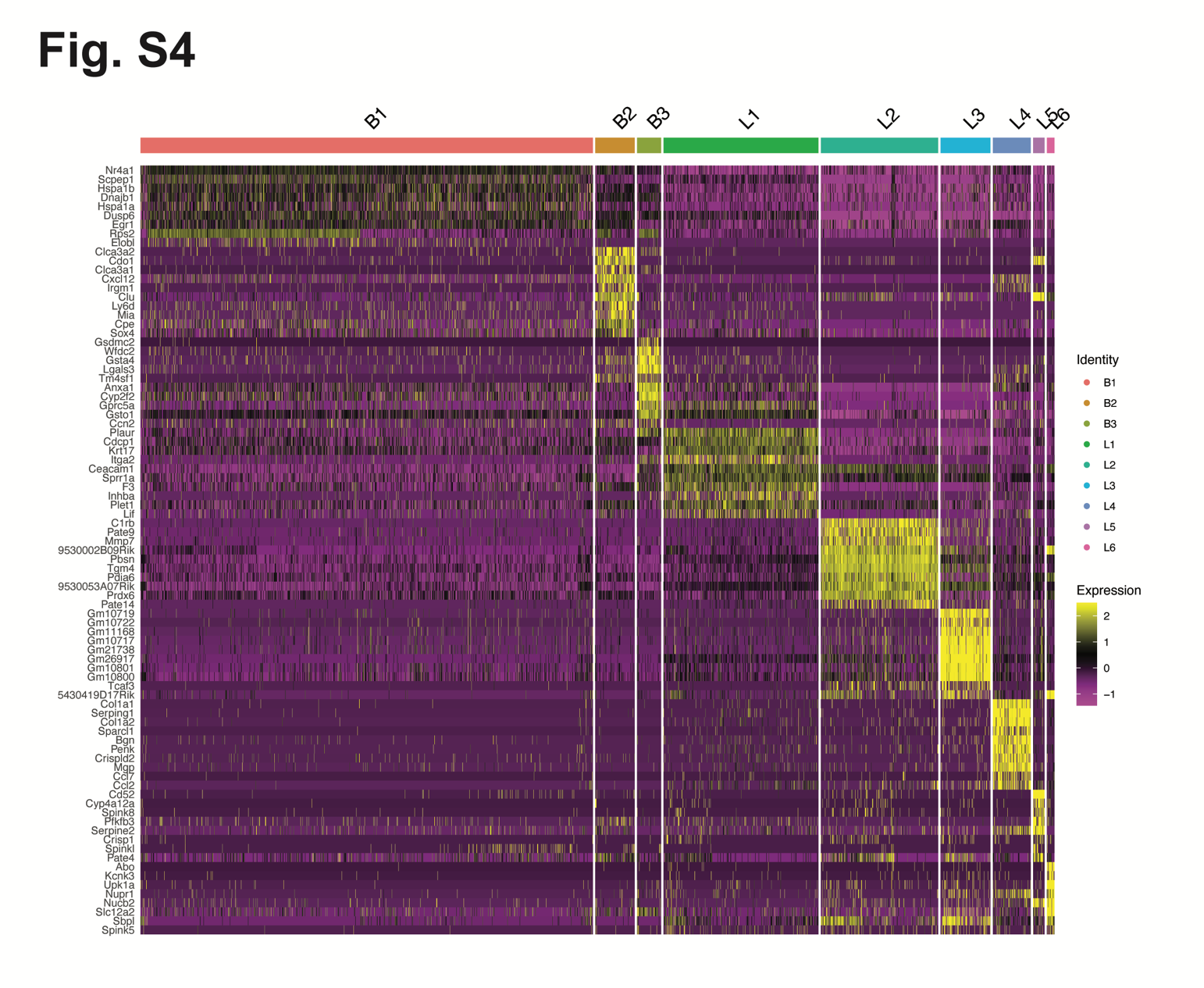

**Fig. S4. Top ten differentially expressed genes in scRNA-Seq clusters.** Heatmap visualization of hierarchical clustering of 10 most differentially expressed genes per group of major cell types identified from GEMMs. B1, B2, and B3 are clusters from the CK5-eYFP basal model. L1 to L6 are clusters with origin from the luminal model.

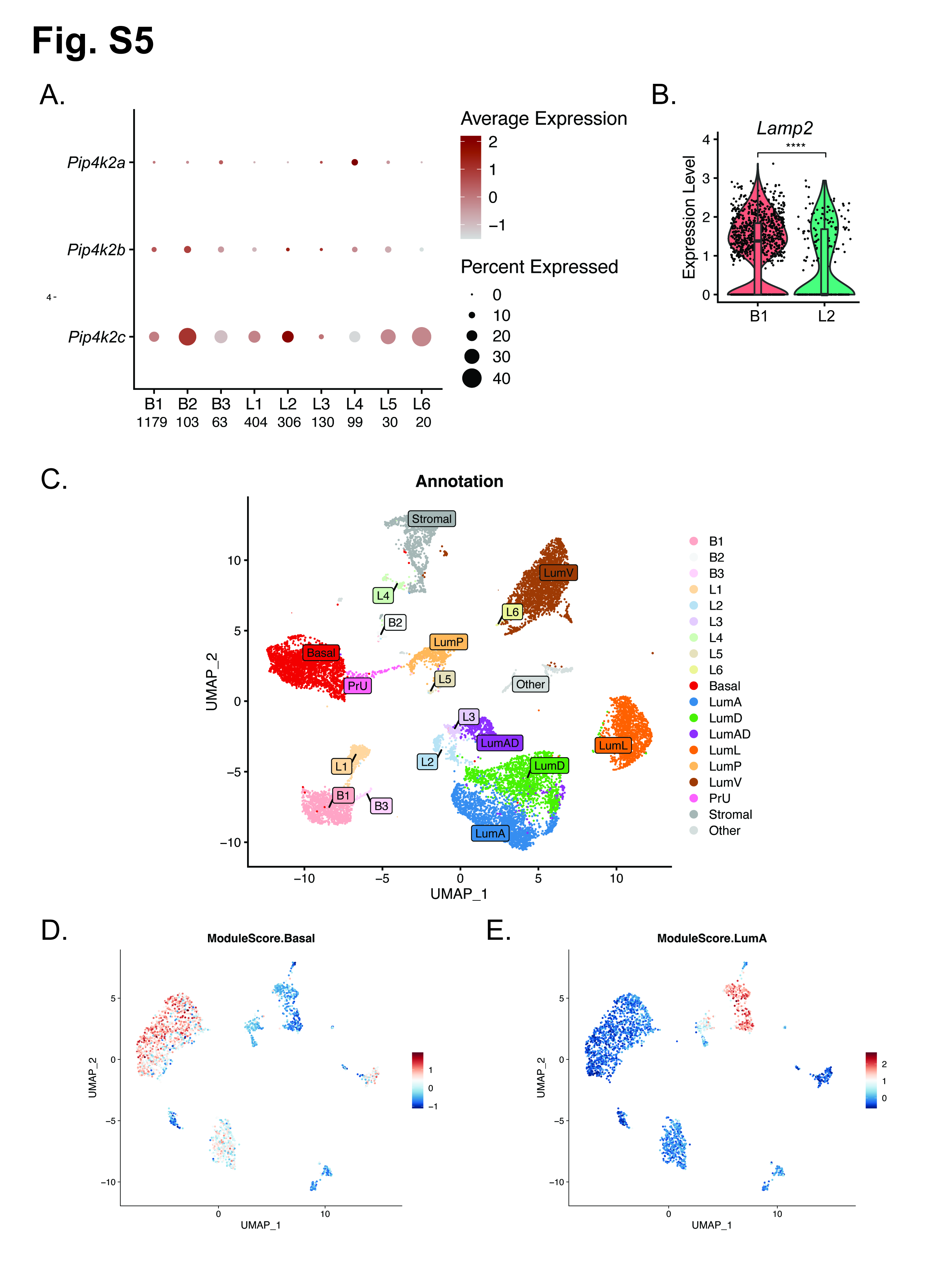

**Fig. S5. scRNA-Seq comparison with atlas datasets.** (**A**) Dot blot indicating relative population and estimated abundance of *Pip4k2* isoform expression in normal adult mouse prostate based on scRNA-Seq clusters identified from basal and luminal GEMMs. As well, (**B**) strong enrichment of lysosome marker *Lamp2* is detected in L2 luminal cluster (LumA) relative to the B1 basal clusters (-0.50 log2FC, adj pval = 1.13E-06). (**C**) UMAP of merged datasets with whole mouse prostate annotations from Crowley *et al. 2020*. Dimly shaded clusters L1-6 and B1-3 from this study, darker spots indicated other atlas data. Modules scores generated from published gene list used to highlight cell type cluster identities of (**D**) basal cells and (**E**) anterior prostate luminal cells (LumA).

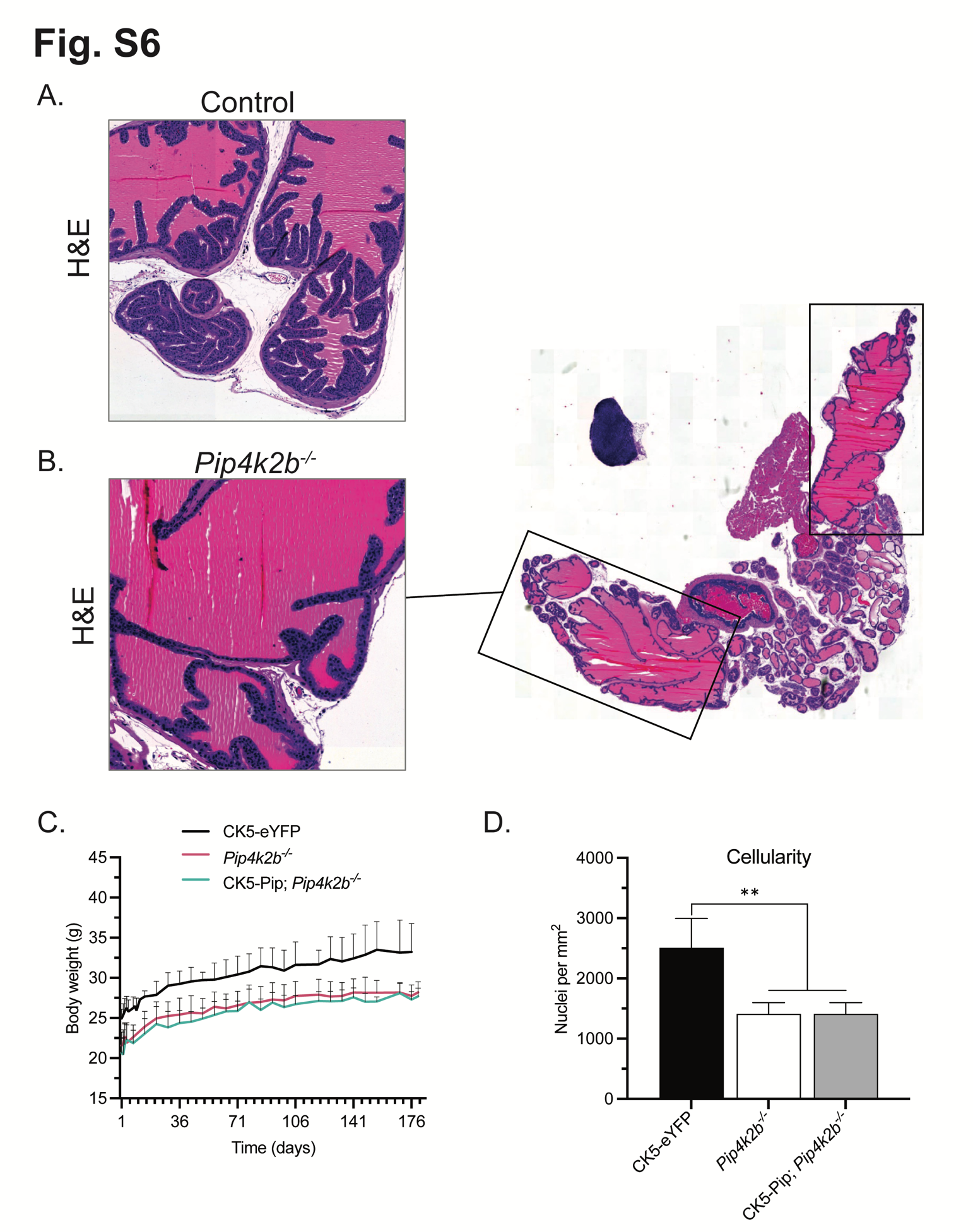

**Fig. S6. Prostate morphology resulting from genomic loss of *Pip4k2b* isoform*.*** At 6-month timepoint, the morphology of mouse prostate lobes is shown for (**A**) eYFP control and (**B**) *Pip4k2b^-/-^* animals. (**C**) Animal body weight and (**D**) anterior prostate lobe cellularity (0.56 fold, pval = 3.4E-03) is summarized. *t* test values: n.s., not significant (*p*> 0.05), * *p* <0.05, ** *p* <0.01, *** *p* < 0.001.

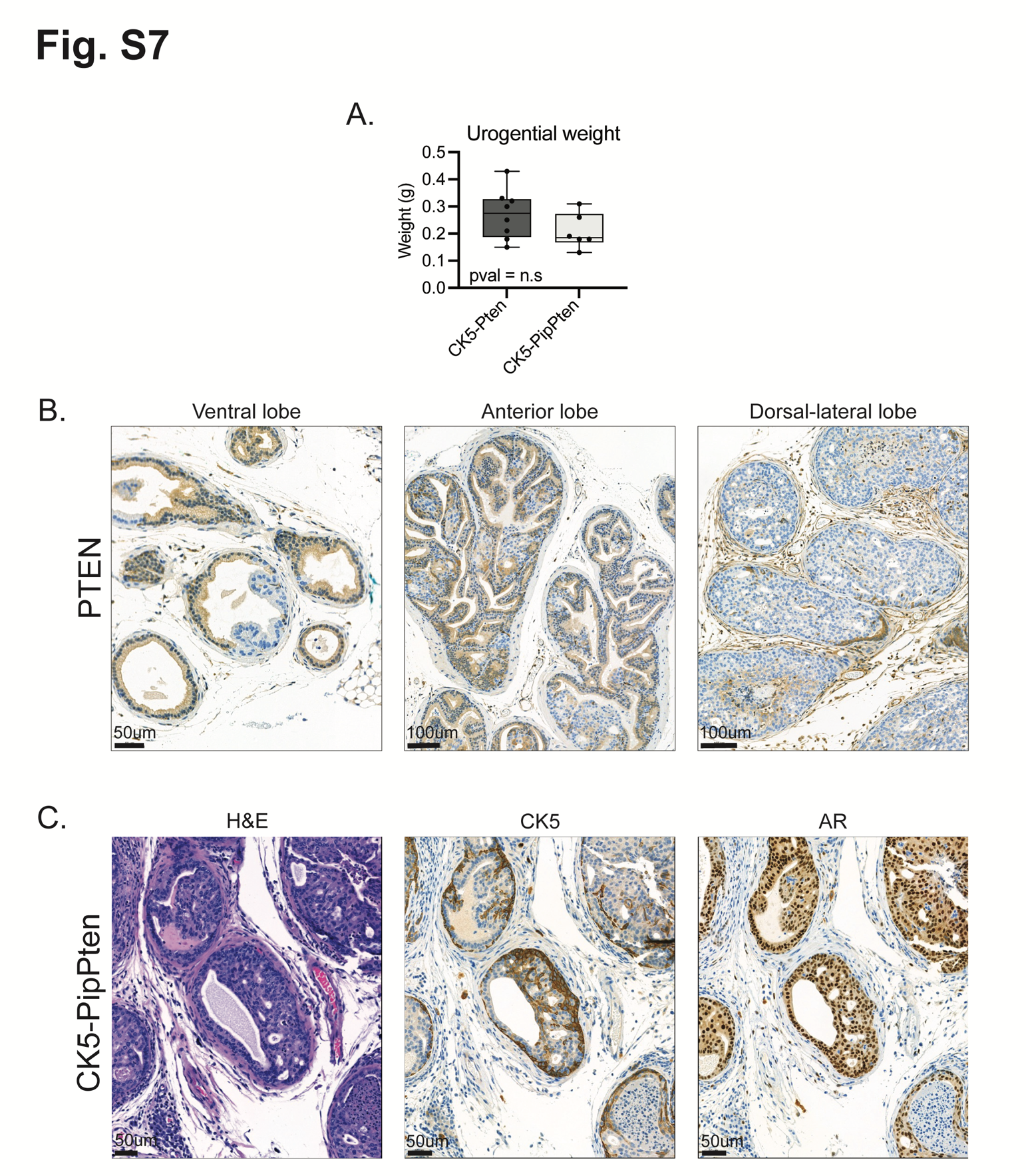

**Fig. S7. Immunohistochemical staining of CK5-PipPten model.** CK5-PipPten prostate tissue from the 3-month timepoint after Cre activation was evaluated. (**A**) Weight of mouse urogenital tracts at time of necropsy are compared for CK5-Pten and CK5-PipPten animals (pval = n. s). (**B**) Immunohistochemical staining for PTEN protein confirms activation of Cre genetic deletion in all prostate lobes. (**C**) Example IHC detection demonstrating basal cell hyperplasia and neoplastic luminal differentiation of prostate glands from CK5 and AR, respectively.

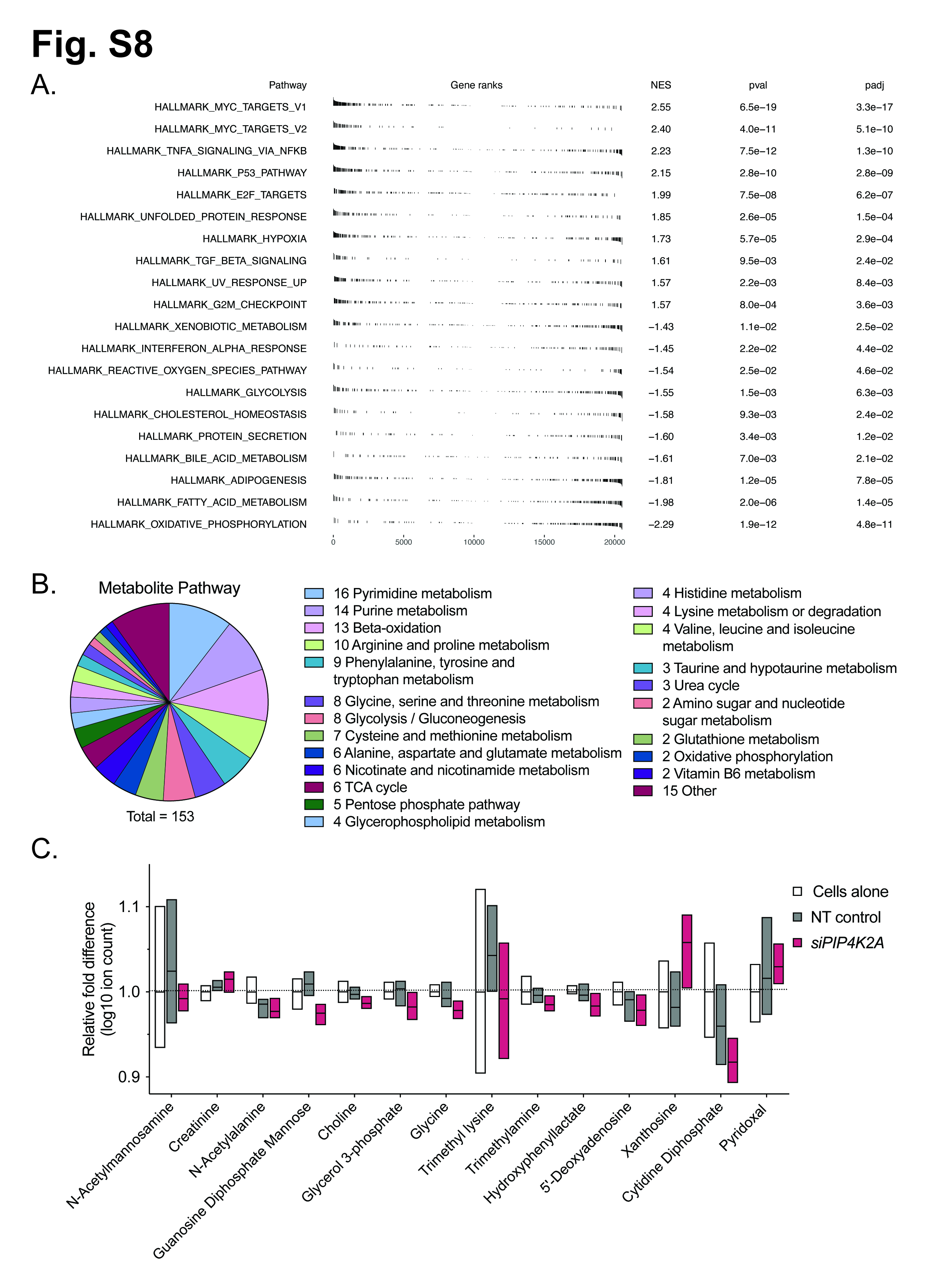

**Fig. S8. Differentially abundant pathways and metabolites from targeting *PIP4K2A*.** (**A**) Normalized enrichment score (NES) from Hallmark gene sets enriched in CK5-Pip eYFP positive cells compared to CK5-eYFP using scRNA-Seq. Levels of significance associated with **Fig. 6.** Polar metabolite abundance was measured in LNCaP cells cultured in androgen-deprived medium and treated with non-targeting RNA or siRNA targeting *PIP4K2A* for 48hrs. (**B**) A total of 153 metabolites were measured an annotated thatr represent multiple energy pathways. (**C**) The most significantly differing metabolites, in exclusion of metabolites shown in **Fig. 6.**, are shown. Each treatment group represents n = 6 biological replicates. See **table S5** for complete data values.

**Supplemental Tables**

**Supplemental Table S1. Summary of genetically engineered mouse models (GEMMs)**

| **Designation** | **Species** | **Genotype** | **Source or Reference** | **Additional Information** |
| --- | --- | --- | --- | --- |
| C57BL/6J | *Mus musculus* | Wild type | Charles river | Adult animals |
| CK5-CreER^T2^ | *Mus musculus* | B6N.129S6(Cg)-*Krt5^tm1.1(cre/ERT2)Blh^*/J | Jackson Laboratory #029155 |  |
| R26R-eYFP | *Mus musculus* | B6.129X1-Gt(ROSA)26Sor*^tm1(EYFP)Cos^*/J | Jackson Laboratory #006148 | Shared by Marianna Kruithof-de Julio lab |
| *Pip4k2a^flox^* | *Mus musculus* | B6;129S-Pip4k2a<tm1.1Lca> | Lewis Cantley lab |  |
| *Pip4k2b^-/-^* | *Mus musculus* | B6;129S-Pip4k2b<Gt(Betageo)1Lca> | Lewis Cantley lab |  |
| *Pten^flox^* | *Mus musculus* | B6.129S4-*Pten^tm1Hwu^*/J | Jackson Laboratory #006440 |  |
| PB-eYFP | *Mus musculus* | B6.DBA-Tg(Pbsn-cre)4Prb x  B6.129X1-Gt(ROSA)26Sor<tm1(EYFP)Cos>/J | *Triscott et al. 2023* |  |

**Supplemental Table S2. Antibodies and reagents used**

| **Antibody or Reagent** | **Company** | **Catalog No.** | **Purpose** | **Conditions** |
| --- | --- | --- | --- | --- |
| PIP4K2A (D83C1) Rabbit mAb | Cell Signaling | 5527S | Western blot | 1:1000 in 5% Milk |
| Anti-PIP4K2A Rabbit Polyclonal Antibody | ProteinTech | 12469-1-AP | IHC | 1:200, HIER TRIS |
| Beta-Actin [AC-15] | Abcam | Ab6276 | Western blot | 1:5000 in 5% BSA |
| PTEN | Cascade BioScience | ABM-2052 | IHC | 1:200, HIER Citrate |
| Ar | Dako‐Agilent | M3562 | IHC | 1:100, HIER TRIS |
| GAPDH | Cell Siganling | 2118 | Western blot | 1:2000 in 5% BSA |
| Keratin 5 polyclonal (CK5) | BioLegend | 905901 | IHC and western blot | 1:5000, HIER TRIS |
| GFP Antibody [DyLight 488] | Novus Biologicals | NBP1-69969 | IHC | 1:200, HIER TRIS |
| Alexa Fluor® 488-AffiniPure Rabbit Anti-Goat IgG (H+L) | Jackson ImmunoResearch | 305-545-003 | IF | 1:200 in Goat serum |
| Cy3-AffiniPure Goat Anti-Rabbit IgG (H+L) | Jackson ImmunoResearch | 111-165-003 | IF | 1:200 in Goat serum |
| Autofluorescence Quenching kit | Vector TrueVIEW | SP-8500 | IF | 1 minute incubation |
| Corn oil | Sigma-Aldrich | C8267 | *In vivo* |  |
| Tamoxifen ≥99% | Sigma-Aldrich | T5648 | *In vivo* | 75mg/kg (daily for 3x) |
| Fetal Bovine Serum, charcoal stripped | Thermo Fisher Scientific | A3382101 | *In vitro* androgen depletion | 5% in Light RPMI |
| ON-TARGETplus Human PIP4K2A (5305) siRNA - SMARTpool | Dharmacon | L-006778-00-0005 | *In vitro* gene knock down | See *Triscott et al. 2023* |
| ON-TARGETplus Non-targeting Pool | Dharmacon | D-001810-10-05 | *In vitro* gene knock down | See *Triscott et al. 2023* |

**Table S3. Crowley et al markers for scRNA-Seq module scores (Excel file)**

**Table S4. GSEA pathways from Fig. 6 (Excel file)**

**Table S5. Metabolomics summary (Excel file)**
